# Brain structure, phenotypic and genetic correlates of reading abilities

**DOI:** 10.1101/2022.02.24.481767

**Authors:** Amaia Carrión-Castillo, Pedro M. Paz-Alonso, Manuel Carreiras

## Abstract

Reading is an evolutionary new development that recruits and tunes brain circuitry connecting visual- and language-processing regions. We investigated the structural correlates of reading and whether genetics influence brain-reading associations. First, we identified left hemisphere cortical surface area (CSA) and cortical thickness (CT) correlates of reading in the large ABCD dataset (N=9,013) of 9-to-10-year-olds. Next, the heritability of cognitive and brain measures of interest was examined through complementary approaches. Last, shared genetic effects between reading, reading-related cognitive traits and reading-associated brain measures were examined by computing genetic correlations and polygenic score analyses, and through mediation analyses. Our results support that morphometric brain measures are related to reading abilities, and that the total left CSA in general, and left superior temporal gyrus CSA in particular, contribute to reading partially through genetic factors.

## Introduction

Reading requires a brain system capable of integrating orthographic, phonological, and lexico-semantic features of written words (Sandak et al., 2004). The invention of reading approximately 5000 years ago does not provide enough time in the evolutionary scale to develop such a specific circuitry. Thus, reading recruits already available networks in the brain implicated in language and visual processing (Dehaene & Cohen, 2007), including the inferior frontal gyrus (IFG), the superior temporal gyrus (STG), the inferior parietal lobe (IPL), and occipito-temporal regions (fusiform gyrus, inferior temporal gyrus; e.g., Friederici, 2012; Lau et al., 2008). The development of this reading network, which includes a dorsal and a ventral processing stream (Pugh 2001, Martin et al., 2015), is shaped by the literacy environment and genetic constraints.

Genetic variation explains a substantial component of reading abilities, with twin-based heritability (twin-h^2^) estimates of 0.66 (Andreola et al., 2021) and population-based heritability (SNP-h^2^) estimates of 0.50 (Verhoef et al., 2020). The largest genome-wide association study (GWAS) of language and reading-related traits to date (N~34,000) has confirmed the robust heritability estimates for these traits, identifying a single genome-wide significant locus for word reading in chromosome 1 (Eising et al., 2021). This study has also highlighted a shared genetic component of reading-related measures with other cognitive components and the cortical surface area (CSA) of the banks of the left superior temporal sulcus (STS; Eising et al., 2021). Dyslexia also has a complex genetic and environmental aetiology, with twin-h^2^ estimates of 0.40-0.60 (Fisher & DeFries, 2002) and SNP-h^2^ estimates of 0.15-0.19 (Doust et al., 2021; Gialluisi et al., 2020). Another recent GWAS study with an unprecedented sample size identified 42 loci associated with dyslexia at the genome-wide significant level, consistent with high polygenicity of the trait (Doust et al., 2021). Recently, the brain imaging genetics field has also been revolutionized by meta-analytic efforts (Grasby et al., 2020) and large-scale datasets such as the UK Biobank (UKB; Elliott et al., 2018; Smith et al., 2021), providing new insights into the role genetics play shaping brain structure. However, a mechanistic account of how these genetic effects contribute to the neurobiology of cognitive functions and human behaviour is still lacking.

Over the last decade, there has been an increase of studies on the brain imaging genetics of reading, examining the association between a wide range of neuroimaging phenotypes and mostly candidate genes for reading (see Landi & Perdue, 2019 for a review), as well as a few genome-wide studies (e.g., Roeske et al., 2009; N=200). Functional studies so far have relied on small samples (range: 33-427 participants; Landi & Perdue, 2019) and have shown mixed findings, reflecting the challenging task of characterizing the reading phenotype and combining it with informative task designs, as well as the difficulty to transfer those functional designs into larger datasets. Although structural imaging phenotypes have been studied in slightly larger sample sizes (range: 56-1,717; Landi & Perdue, 2019), these are still far from the sample sizes required to identify robust associations given the small genetic effects that are expected, given the polygenicity of both reading (Eising et al., 2021) and brain phenotypes (Smith et al., 2021). Hence, these studies provide a groundwork to understand the link from genetic variation to brain phenotypes and reading, but systematicity in the analytical approaches is lacking as, until now, only a handful of phenotypes and genetic loci have been considered.

It is critical to use large datasets to identify robust and scalable brain correlates of reading to perform genetic analyses and to seek for the replicability of the results. The goal of the present study was two-fold: (A) to identify robust brain correlates of reading in the developing brain; and (B) to examine whether genetics influence brain-reading associations (see Figure 1). To address these goals we used three complementary analytical approaches, namely (1) conducting a regression analysis in the large Adolescent Brain Cognitive Development (ABCD) dataset (N=9,013) to define left hemisphere structural cortical measures robustly associated with reading performance (goal A; Figure 1); (2) examining the genetic architecture of reading-related cognitive and brain measures by estimating the heritability of these traits using the ABCD and other publicly available datasets (goal B); and (3) exploring the shared genetic influences on brain and reading through genetic correlation and polygenic scoring combined with mediation analyses (goal B).

**Figure 1.**
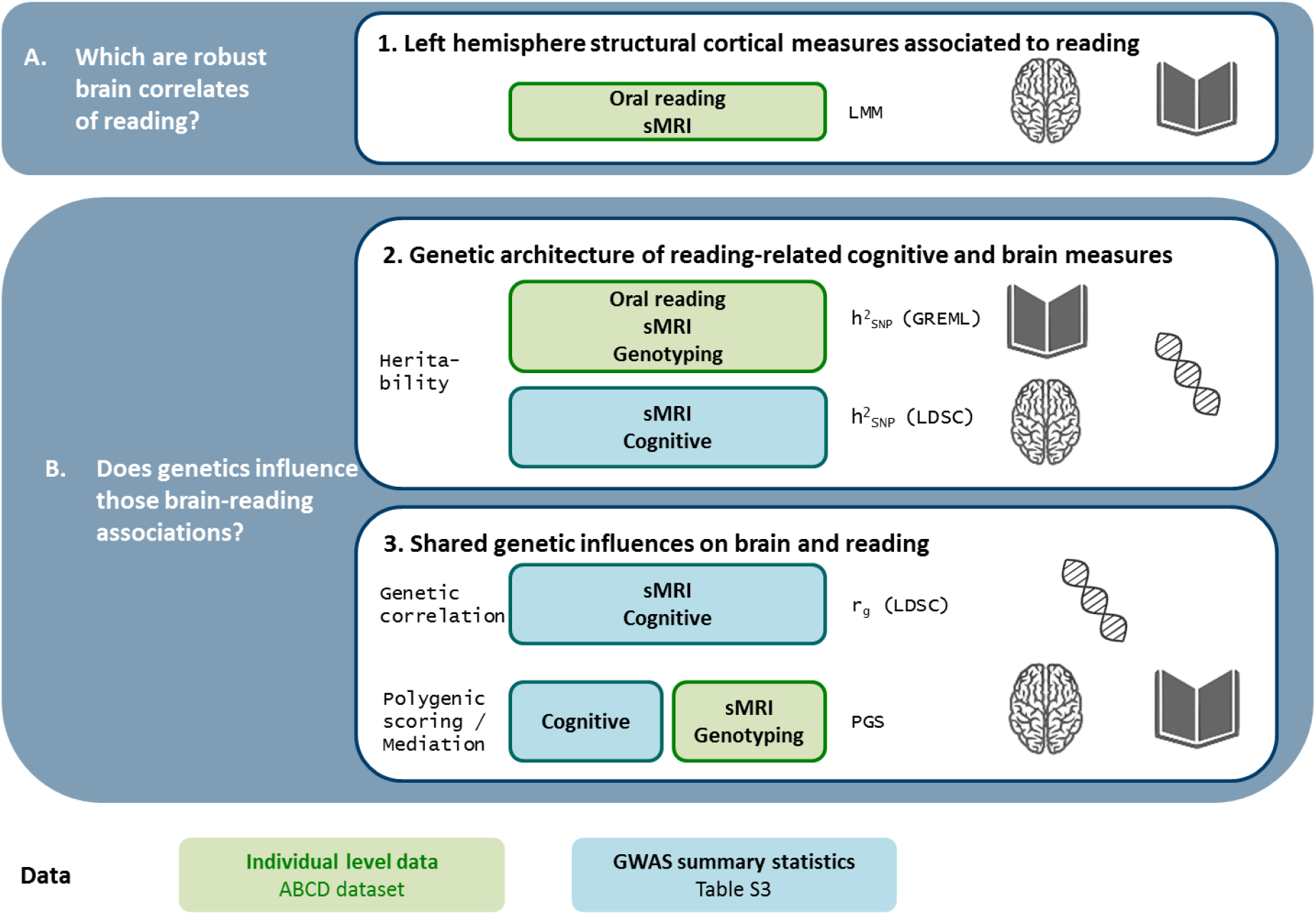
Overview of study goals and analytical approaches. A and B state the main goals of the study, and the analytical steps taken to address them are depicted in the central panels (1, 2, 3). The measures used in each analysis are specified in these panels. The data used for each analysis type are indicated by the colour: green (individual level data from the ABCD study) and blue (GWAS summary statistics). sMRI= structural MRI measures of cortical thickness (CT) and cortical surface area (CSA); LMM= linear mixed effect model; h^2^_SNP_(GREML)= SNP-heritability estimated from unrelated samples using the genome-based restricted maximum likelihood (GREML) estimation approach; h^2^_SNP_ (LDSC)= SNP-heritability estimated from GWAS summary statistics using linkage disequilibrium score regression (LDSC); PGS= polygenic scoring analysis.

In sum, we expect to reliably identify brain-behaviour associations within the known reading network, and to take advantage of the large ABCD dataset to unravel other more subtle but robust associations that have not been detected in smaller datasets (Dick et al., 2021). As speech comprehension ontogenetically precedes literacy (Rueckl et al., 2015), we hypothesize that hub regions of the speech processing network such as the IFG or STG will be associated with reading abilities. Moreover, if genetic effects mediate this effect, they are likely to act upon those regions, more than in other regions such as the ventral occipito-temporal cortex, which is a plastic and environmentally malleable area (Dehaene et al., 2015).

## Results

### Left hemisphere structural correlates of reading abilities

To identify robust cortical structural correlates of reading in the developing brain (Figure 1A) we performed a regression analysis using the ABCD study dataset (N=9,013, Figure S1). Variable definitions and sescriptive statistics are shown in Tables S1 and S2. In a baseline model we assessed the effect of covariates (Equation 1), which indicates that age, socioeconomic factors and the first four genetic components are associated with reading (Table S3, Figure S2), while sex is not. We next tested the effect of 150 left-hemisphere measures on reading: the global measures of total CSA and mean CT, and 74 regional measures for CSA and CT using the Destrieux parcellation. We report the effect prior (Model 1) and after (Model 2) adjustment for global brain measures for the regional measures (Table S4). The global measures included in Model 2 were mean left-hemisphere CT for regional CT measures and total left-hemisphere CSA for regional CSA measures.

The strongest association was observed with total CSA (χ^2^(1)=115.11; T=10.75, q-value=1.27e-24). This was reflected in 69 out of 74 of the regional CSA measures being associated with reading in Model 1 (Figure 2A, Table S4), although the majority of these (67/69) were no longer significant after adjusting for total CSA (Model 2), suggesting that in those regions the association with reading abilities was driven by total CSA. Nevertheless, two regional CSA measures were significantly associated with reading in both models: the CSA of the lateral STG, which was consistently positively associated with reading (Model 1: χ^2^(1)=96.14; T=9.82, q-value=8.99e-21 and Model 2: χ^2^(1)=16.65; T=4.08, q-value=0.0017; Figure 2A,C), and the CSA of the superior parietal gyrus, which showed a positive relationship to reading that was shifted to negative after adjustment (Model 1: χ^2^(1)=9.48; T=3.07, q-value=0.0047 and Model 2: χ^2^(1)=9.63; T=-3.1, q-value=0.036). This reversal in the direction of effect reflects that the relative size of CSA in this region is negatively associated with reading abilities, supporting the sensitivity approach to assess the effect of each measure (Hyatt et al. 2020; Dick et al. 2021).

**Figure 2.**
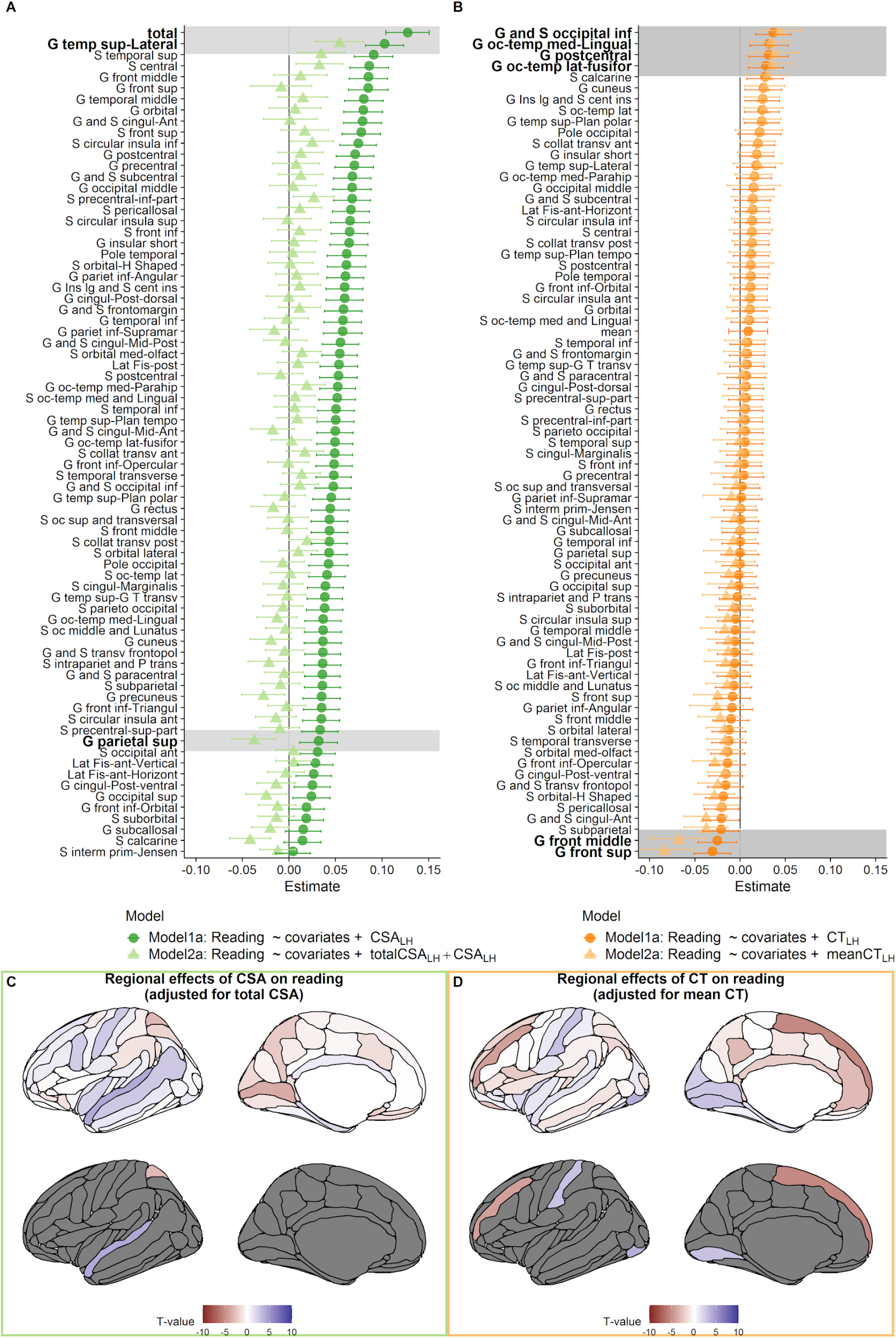
Effect of left-hemisphere cortical measures on reading abilities. Beta estimates for **A** cortical surface area measures and **B** cortical thickness measures. Covariates: ***sex*** + ***age*** + ***high.educ*** + ***household. income*** + ***PC*1:*PC*10**; Model 1: test model, Model 2: test model after adjusting for global measure (total CSA or mean CT). The regions that survived multiple comparisons correction are shaded in grey. **C**, **D**: brainplots of T-values associated for each brain region in Model 2. The upper panels show T-values for all regions, and the lower panels show only T-values for regions that survive multiple comparisons correction. CSA= cortical surface area; CT= cortical thickness; LH= left-hemisphere; G = gyrus; S = sulcus.

CT of six regions was robustly associated with reading abilities (Figure 2B, Table S4): the postcentral gyrus and three occipital regions (occipital inferior gyrus and sulcus, occipito-temporal medial lingual gyrus, occipito-temporal lateral fusiform gyrus) were positively and robustly associated with reading, while two superior frontal regions (middle frontal gyrus, superior frontal gyrus) showed a negative association. This negative relationship increased when adjusting for mean CT (Model 2), again suggesting a possible paradox.

The specificity of the observed associations was assessed through sensitivity analysis by adjusting for additional cognitive variables that were correlated with reading abilities (Figure S3, Table S5), namely fluid intelligence, matrix reasoning and/or picture vocabulary. Total CSA, CSA of the lateral STG, CT of the occipital inferior gyrus and sulcus and CT of the occipito-temporal lateral fusiform gyrus were significantly associated with reading abilities across all of these analyses, while the associations were weaker and no longer significant for the other five measures after adjusting for some of the additional cognitive measures, suggesting that the association we observed with reading may be part of a more general cognitive effect in those specific regions (see Table S5, Figure S4).

To establish whether the observed associations were left-hemisphere specific, we also explored the effect of homotopic right-hemisphere measures on reading, which were imperfectly correlated with the left-hemisphere measures (Pearson’s r ranging from 0.38 to 0.8 for the regional measures, see Figure S5). The right total CSA was associated with reading (χ^2^(1)=109.924; T=10.50, p-value=1.16e-25) to a similar extent as the left CSA. At the regional level, the association was robust to adjustment for global measures (Model 2) for three measures (see Figure S6). The CT of the right occipito-temporal lateral fusiform gyrus was consistently associated with reading abilities (Model 1: χ^2^(1)=8.93; T=2.99, p-value=0.0028 and Model 2: χ^2^(1)=8.10; T=2.84, p-value=0.0045), while for the other two measures the direction of the associations reverses when adjusting for global measures: the CSA of the right superior parietal gyrus reversed the direction of effect after adjustment for total right CSA (Model 1: χ^2^(1)=8.47; T=2.9, p-value=0.0036 and Model 2: χ^2^(1)=5.05; T=-2.24, p-value=0.024) and the CT of the right superior frontal gyrus was negatively associated in both models and the association became stronger after adjustment for mean right CT (Model 1: χ^2^(1)=10.01; T=-3.15, p-value=0.0015 and Model 2: χ^2^(1)=35.57; T=-5.97, p-value=2e-9).

Correlations among these nine reading-associated measures showed two independent clusters reflecting the type of measurement: one for the CSA measures and the other for CT measures (Figure S5A). After correction for global measures, the correlations showed an overall decrease in the strength of the effect, indicating that they are mostly independent to each other and therefore not likely to reflect the same relationship with reading abilities (see Figure S5B). After this adjustment, the direction of some correlations was reversed, which may reflect the complex relationship between the regional and global measures: e.g. a weak negative correlation between the CSA of the lateral STG and the CSA of the superior parietal gyrus, and weak negative correlations between the CT of the superior and frontal gyri with the rest of the CT measures.

In sum, a set of nine cortical structural measures are robustly associated with reading abilities in this sample. Sensitivity analyses support that several of these effects are robust to other cognitive variables and that the associations are mostly left-hemisphere specific.

### Genetic architecture of reading-related cognitive and brain measures

As a first step to investigate whether genetics contributed to the identified brain-behavioural associations, we established whether our traits of interest were heritable (Figure 1B.2). Heritability was estimated for reading, related cognitive measures and reading-associated brain measures (as defined in Tables S2 and S3) using different methods and datasets: in the ABCD dataset SNP-h^2^ was estimated using genome-based restricted maximum likelihood (GREML; Yang et al., 2010) in a set of unrelated individuals and SNP-h^2^ were computed through linkage-disequilibrium score regression (LDSC; Bulik-Sullivan et al., 2015) from summary statistics of GWASes from publicly available datasets (see Table S6). The heritability measures across traits and methods are reported in Table S7 and Figure S7.

Reading had a SNP-h^2^ of 0.14 in the ABCD dataset, which was nominally significantly different from 0 (p-value=0.025), while the remaining cognitive measures were low to moderately heritable, ranging from 0.09 for WISC-V (p-value=0.09) to 0.25 for vocabulary (p-value=4.6e-4). All nine brain structural measures associated with reading were also heritable (Figure S7B), with the CT of the postcentral gyrus having the highest estimate (SNP-h^2^=0.32, p-value=1.07e-6), and the inferior occipital gyrus and sulcus having the lowest (SNP-h^2^=0.11, p-value=0.02). All were nominally significantly different from zero, although four (two regional CSA and two occipital CT measures) did not survive the correction for the nine tested measures. After the adjustment for global brain measures, most regional estimates were lower (Table S5; Figure S7), and only two CT measures were nominally (CT of middle frontal gyrus) or significantly (CT of the postcentral gyrus) different from zero.

SNP-h^2^ estimates from summary statistics from GWASes were moderate for dyslexia, educational attainment (EA) and cognitive performance, consistent with original GWAS reports (Table S7; Gialluisi et al., 2020; Lee et al., 2018). All of the SNP-h^2^ (LDSC) estimates for brain measures, derived from the UKB dataset, were significantly different from zero (p-values<0.05/12; see Table S7), ranging from 0.31 (total CSA) to 0.12 (CT of the occipital inferior gyrus and sulcus).

In sum, these analyses support that reading and reading-associated cognitive and brain measures have a genetic component.

### Shared genetic influences on brain and reading

Having established that genetic variation explains part of the variance of the reading-associated brain measures, we explored the extent to which genetics mediate brain-behaviour associations (Figure 1B.3). To this end, we first computed genetic correlation estimates. Then, we followed up the most promising brain measures with polygenic score (PGS) analyses. Finally, we conducted a mediation analysis to examine whether CSA measures mediate PGS effects of cognitive performance on reading.

#### Genetic correlation between cognition and reading-associated brain measures

Genetic correlation (r_g_) is the proportion of variance that two traits share due to genetic variance. We estimated genetic correlations across cognitive traits and brain measures using LDSC. Bivariate GREML analysis was not considered as it was not well-powered to detect genetic correlations between brain measures and reading in the ABCD dataset (see Figure S8).

Genetic correlations (Figure 3) revealed a low positive correlation of total CSA with EA (rg=0.051, p-value=0.0365) and cognitive performance (r_g_=0.0741, p-value=0.0105). Among the regional measures, the CSA of the lateral STG was the only measure showing positive genetic correlations with EA (r_g_=0.056, p-value=0.0379) and cognitive performance (r_g_=0.075, p-value=0.0277), and a negative genetic correlation with dyslexia (r_g_=-0.314, p-value=0.0207). Although these genetic correlations were nominally significant they did not survive multiple comparison corrections. However, they all showed a consistent direction of the effect indicating a shared genetic overlap between reading and related cognitive measures and CSA measures (i.e., total, lateral STG). These two measure were further analyzed using PGS.

**Figure 3.**
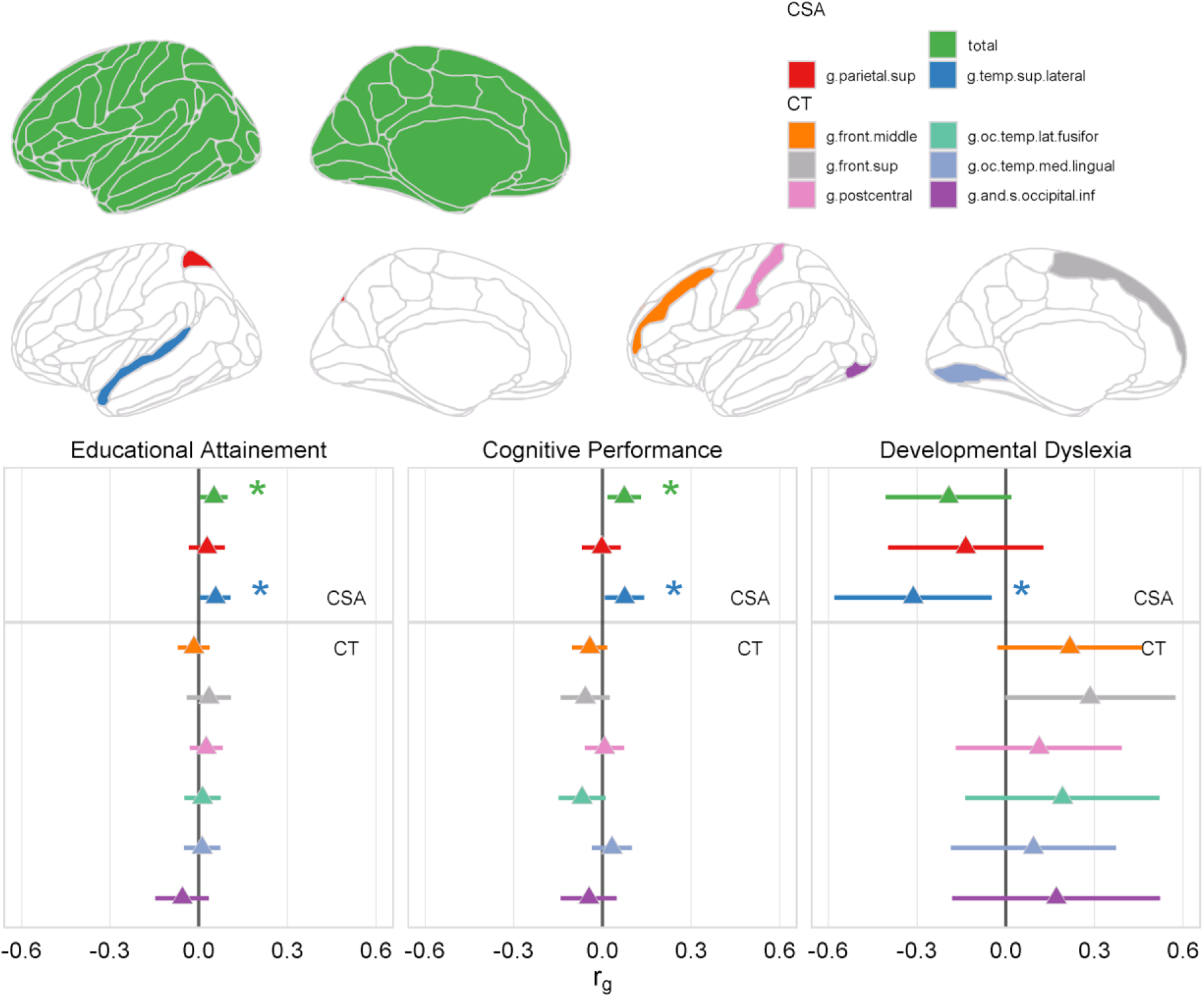
Genetic correlation (r_g_) between the reading-associated left-hemisphere brain measures and cognitive traits (method LDSC). * = p-value <0.05. LDSC = linkage disequilibrium score regression. CSA = cortical surface area; CT = cortical thickness; G = gyrus; S = sulcus; front. = frontal; oc. = occipital; temp.= tempora; sup = superior; inf = inferior.

#### Polygenic scores of cognition predict reading and reading-associated brain measures

PGS are individual level predictors derived from the sum of effect alleles at a SNP, weighted by the regression coefficient describing each SNP’s level of association with a trait. PGSes can also be used to study the genetic relationship between two traits by making predictions across traits (Maier et al. 2018).

We first used GWASes for EA and cognitive performance to define the PGS that best predicted reading in the ABCD dataset. The PGS for cognitive performance (PGS_CP_) at the p-value threshold of 0.05 explained the largest amount of variance in reading abilities (*ΔR*^2^=3.6%, p-value=5.5e-38;Figures S7, S8). Therefore, we used this best PGS to perform cross-trait prediction in brain measures.

This PGS_CP_ was a significant predictor of the left CSA brain measures associated with reading, explaining up to 0.75% of the variance of the total CSA (p-value=1.3e-11), and up to 0.56% of the CSA of the lateral STG (p-value=1e-07). These results indicate that genetic effects that contribute to cognitive performance also influence CSA measures associated with reading. The regional prediction was significantly diminished and no longer significant when including the total CSA as a covariate in the regression analysis (*ΔR*^2^=8.35e-05, p-value=0.176), suggesting that the genetic effect is shared between the total CSA and the CSA of the lateral STG (Figure 4).

**Figure 4.**
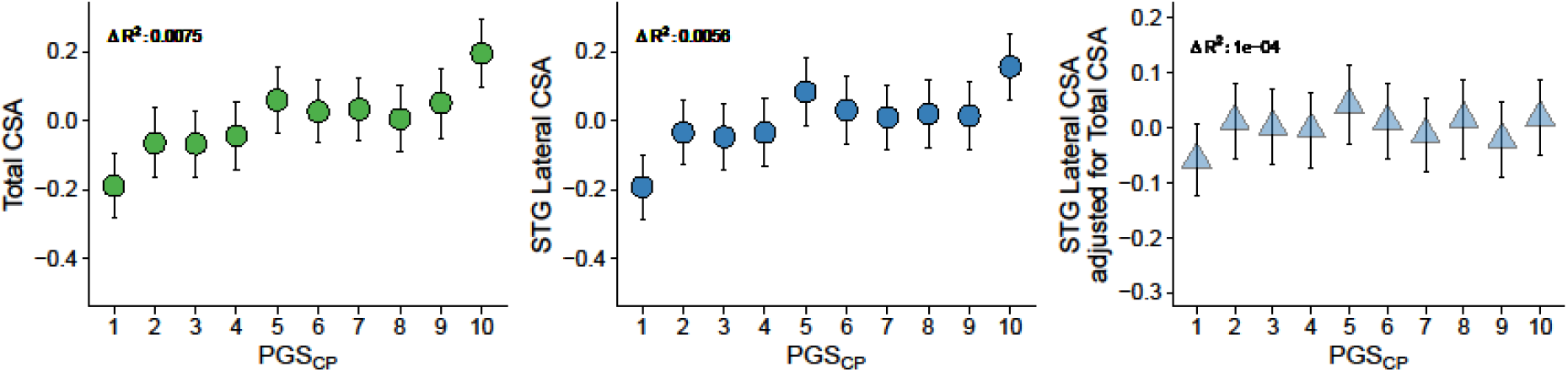
Decile plots for PGS of cognitive performance on left-hemisphere CSA measures. The points indicate the mean CSA for each decile, the error bars show the 95% confidence intervals. PGS_CP_= polygenic score for cognitive performance at the p-value threshold of 0.05; CSA= Cortical surface area; STG: superior temporal gyrus; *ΔR*^2^: variance explained by the PGS.

#### The relation between CSA and reading is mediated by polygenic scores of cognitive performance

The relation between reading abilities, total CSA, the CSA of the lateral STG and PGS of cognitive performance was further explored through a set of mediation analyses that are summarized in Table 1. The effect of either CSA measure (total and STG) on reading was partially mediated by the PGS_CP_, the indirect effect explaining 17.4% of the total brain-behaviour effect for total CSA, and 13.2% for the CSA of the lateral STG (Figure 5).

**Table 1:**
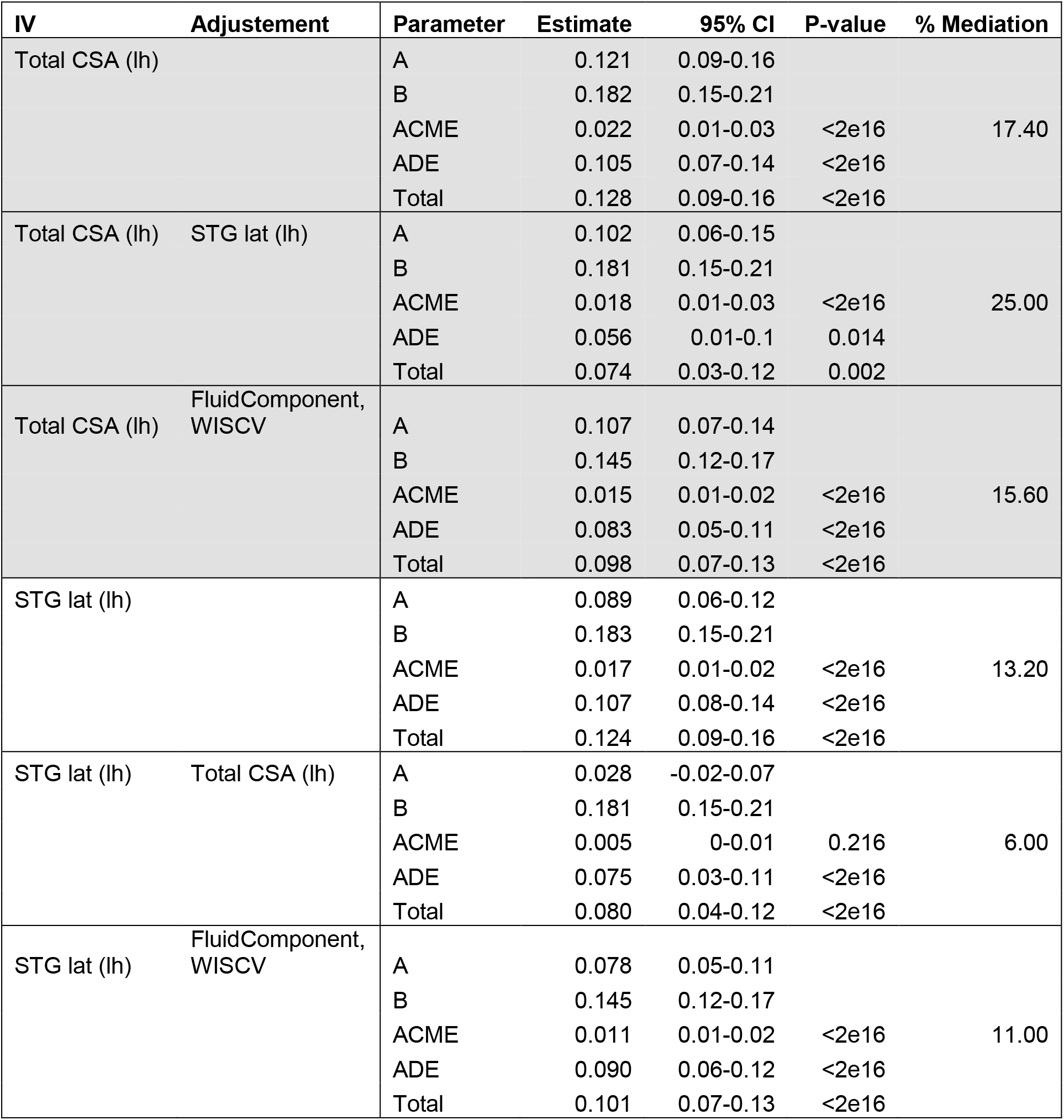
Mediation model testing the significance of PGS_CP_ as mediator of the CSA measures (IV) and reading (dependent variable) association. All models included MRI scanner as a random factor, and the following covariates: sex, age, high.educ, household.income, PC1:PC10. IV: independent variable. Adjustement: additional covariates included in mediation models. Estimate: standardized beta for the parameter. 95% CI: confidence intervals for the estimate. A: Effect of the independent variable on the mediator. B: Effect of the mediator on the dependent variable. ACME= Average Causal Mediation Effect; ADE= Average Direct Effect. PGS_CP_= polygenic score for cognitive performance at the p-value threshold of 0.05; CSA= Cortical surface area; STG lat= lateral part of the superior temporal gyrus. Lh= left-hemisphere. WISCV= WISC-V Matrix Reasoning Total Scaled Score.

**Figure 5.**
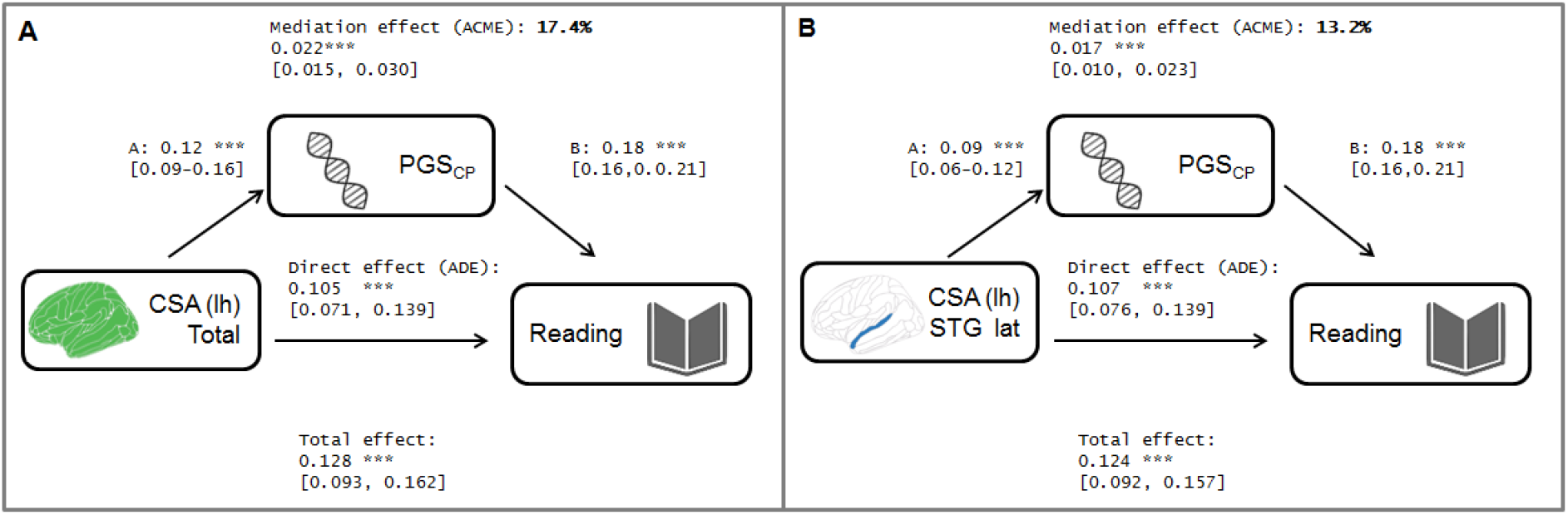
Mediation of the PGS_CP_ on the association between the CSA measures and reading. **(A)** total CSA as the independent variable and **(B)** CSA of the lateral STG as the independent variable. The standardized estimates of the paths are provided, with the 95% confidence intervals in brackets. ACME= Average Causal Mediation Effect; ADE= Average Direct Effect. A: Effect of the independent variable on the mediator. B: Effect of the mediator on the dependent variable. PGS_CP_= polygenic score for cognitive performance at the p-value threshold of 0.05; CSA= Cortical surface area; STG lat= lateral part of the superior temporal gyrus.

When adjusting the CSA measures with each other, the indirect effect was no longer significant, which again supports that the genetic effect is common to both CSA measures (Table 1). For sensitivity, we also repeated the mediation analyses adjusting reading for general intelligence measures, which did not affect the amount of the mediated effect (see Table 1).

In sum, through complementary approaches, we find evidence for a shared genetic component between reading-related cognitive measures and CSA measures associated with reading, namely the total left-hemisphere CSA and the CSA of the lateral STG.

## Discussion

In the current study, we have established a set of morphometric measures associated with reading abilities in the ABCD dataset, constituted by 3 CSA and 6 CT measures, including relevant regions of the reading network. Next, we explored if genes played a role in the brain-behaviour relationship and found evidence for genetic overlap with two CSA measures (total CSA and CSA of the lateral STG) but not for any of the CT measures. Finally, through mediation analysis, we show that the effect of the CSA measures on reading is mediated through PGS on cognitive performance. These results are discussed next.

### Cortical reading network

A set of nine cortical structural measures were robustly associated with reading in the ABCD studyof 9-to-10-year-old US children. The unprecedented scale of this dataset (N>9,000) allowed us to perform an unbiased search for morphometric (CSA and CT) correlates of reading. Total CSA was associated with reading abilities, and this effect was reflected by a global association of most regional CSA measures. A previous study that used the ABCD dataset also demonstrated that performance on crystallised cognitive measures including reading was more strongly associated with total CSA compared to the regional CSA measures (Palmer et al., 2019). Nevertheless, we also observed a regional effect beyond the generalized effect, as the CSA of the lateral part of the STG was associated with reading skills after adjusting for total left CSA. The STG is a known hub of the speech and reading networks (Hickok and Poeppel, 2007; Rueckl et al., 2015): functional MRI studies have shown that the posterior part of the STG is a multimodal area key for audiovisual integration of speech and print, involved in the grapheme-to-phoneme correspondence mapping within the dorsal reading pathway (Atteveldt et al., 2004; Lau et al., 2008; Boets et al., 2013), while the anterior STG seems to be more related to the ventral pathway of speech processing (Rueckl et al., 2015; Friederici et al., 2012). The present study included the CSA of the lateral STG as a single unit, and we cannot, therefore, relate the observed effects to the different specific processes within it. Nevertheless, it is noteworthy that this specific region, the key hub for the ventral and dorsal reading networks, was the most strongly associated with reading abilities. Sensitivity analyses showed that the association was not driven by more general cognitive processes, as it was robust to adjustment of other cognitive variables (i.e. fluid intelligence and matrix reasoning). Recent studies looking at grey matter measures and reading have also highlighted the relevance of the STG. Plonski et al. (2017) performed multivariate classification of children with dyslexia based on morphometric features and identified the mean curvature of the lateral STG as a feature discriminating dyslexics from controls, and Perdue et al. (2020) found that the CT of the left STG was positively correlated with word and pseudoword reading performance.

Six regional CT measures were associated with reading abilities, although mean CT was not. The postcentral gyrus and three occipital CT measures (inferior occipital gyrus and sulcus, lingual gyrus, fusiform gyrus) were positively associated with reading abilities after adjusting for other cognitive covariates. These occipital regions are part of the visual system and are involved in the left ventral occipito-temporal network of reading that performs visual-orthographic processing (e.g., Ben-Shachar et al., 2007), with the left fusiform gyrus including the so-called visual word form area (Cohen et al., 2000). The three other regional measures associated with reading abilities were the CT of the superior and middle frontal gyri and the CSA of the superior parietal gyrus. The left parietal cortex has been implicated in letter position coding (Carreiras et al., 2015), while activation in the middle frontal gyrus is activated by reading in Chinese and French children (Feng et al., 2020). However, the direction of the association of these measures with reading either shifted the direction of effect (for the superior parietal) or the extent of the association became bigger (for frontal CT regions) after controlling for the global measures, implying that these effects are dependent on adjustment to global measures and should be interpreted with caution.

In sum, we established a set of structural correlates of reading abilities, which included key regions such as the CSA of the STG involved in the dorsal and ventral reading networks, and the CT of occipital regions implicated in the ventral reading network. These associations were subtle but robust across sensitivity analyses. No association was found between structural measures of other key regions of the reading network, such as the inferior frontal gyrus or inferior parietal regions, which may indicate that the multifaceted nature of these regions and that their implication in reading is not reflected by morphological changes, at least at the developmental stage (late childhood, 9-10yo) at which we tested this association.

### Genetic architecture of reading and associated measures

Reading was heritable in the ABCD dataset, although with a lower estimate relative to previously reported results for reading in a large GWAS meta-analysis (Eising et al., 2021; Verhoef et al., 2020). Morphometric brain measures associated with reading abilities showed low to modest heritabilities, which were significantly different from zero in most cases, as has been previously reported for these measures (Grasby et al., 2020). Total left CSA was the measure with the highest estimates across methods, while regional measures were more variable. The adjustment of global measures reduced the estimates of most regional measures in the ABCD dataset, many of which were no longer significant. Estimates of SNP-h^2^, based on summary statistics from the UK Biobank dataset, ranged 0.12-0.23, as previously reported (Smith et al., 2021). Note that these brain GWAS summary statistics had been adjusted for multiple imaging confounds, including global measures (i.e. head size; Smith et al., 2021; Alfaro-Almagro et al., 2021). The differences of brain SNP-h^2^ estimates after adjustment for global brain measures in the GREML (ABCD dataset) and LDSC (UK Biobank dataset) could be due to dataset heterogeneity (e.g. age) or may reflect the limited power of the GREML approach in the ABCD dataset (N~4,600; Visscher et al., 2014). Thus, we confirmed that reading and reading-related cognitive and brain traits were moderately heritable across methods.

### Shared genetic effects on reading and CSA measures

The three complementary analyses we performed to further elucidate possible shared genetic effects for reading and reading-associated brain measures included first performing genetic correlation analyses, then examining the most promising signals through polygenic scores before finally assessing these relations through mediation analyses.

There was a consistent trend of a small positive genetic correlation between the total CSA and the CSA of the lateral STG with reading-related cognitive measures (EA and cognitive performance), and a negative genetic correlation with dyslexia, although these associations did not survive multiple comparisons correction. Genetic correlation between total CSA and EA has also been previously reported, albeit with a considerably stronger genetic correlation of 0.22 (p=1.9e-13; Grasby et al., 2020), using the same GWAS summary statistics for EA (Lee et al., 2018) and the ENIGMA GWAS meta-analysis data for total CSA (which includes a subset [N=10,083] of the UK Biobank). To understand the variability between the present and the previous study we computed the genetic correlation of the mean total CSA (ENIGMA) and total left CSA (UK Biobank) measures across datasets (partially overlapping sample). This yielded a genetic correlation of 0.55 (se=0.05), which was significantly different from 1 (p=1e-20). This lack of concordance may be partially explained by the fact that the UK Biobank GWASes we used here were run adjusting for head size (Smith et al., 2021), while the ENIGMA meta-analysis did not correct for it (Grasby et al., 2020). Another recent GWAS meta-analysis of reading- and language-related traits (Eising et al., 2021) found a significant genetic correlation between the reading and the CSA of the banks of the superior temporal sulcus (STS), which partially overlaps the lateral STG we report here. They used the same UK Biobank GWAS resouce used in the present study but a different brain parcellation (Smith et al., 2021) to compute the genetic correlations between the reading and language GWAS meta-analysis and 58 structural neuroimaging traits with known links to reading and language. Hence, our results are in line with the literature, supporting a possible genetic overlap between reading and related cognitive traits with total CSA and the CSA around the superior temporal regions.

We did not find evidence for a genetic overlap between reading and any of the other reading associated CT and CSA measures. The power to detect genetic correlations depends on the strength of the phenotypic association, the strength of the genetic effects on each trait (i.e. heritability) and the actual genetic overlap that exists (Visscher et al., 2014). In our results, the heritability estimates were of a similar order of magnitude for all cortical measures (Figure S7), but the strength of the association with reading was lower for these other measures, which may have resulted in diminished power to detect low to moderate genetic correlations. Note that a lack of a genetic correlation may also occur despite there being a genetic overlap, if there are mixed effect alleles that contribute to both traits (Frei et al., 2019).

To further examine the shared genetic contributions to reading and CSA measures we used polygenic scoring. A polygenic score of cognitive performance (PGS_CP_) predicted 3.6% of the variance in reading abilities and was also a significant predictor of total CSA and the CSA of the STG region. Mediation analyses confirmed that the effect of both of these CSA measures on reading is partially mediated through the PGS_CP_. This mediation effect explains up to 17% of the effect of total CSA, and 13% of the effect of the CSA of the STG, and seems to be shared between the two CSA measures, as the adjustment of the CSA measures with each other diminished or made the mediation disappear. Based on these results, we suggest that part of the effect of the CSA measures is mediated by genetic effects captured by the PGS. This relationship is similar to the one reported for intelligence measures in young-adult samples: CSA measures mediated the effect of the PGS of EA on intelligence in two twin-based datasets (Mitchell et al., 2020) and vertex-wise surface area measures were shown to mediate the effect of PGS of EA on the general intelligence factor ‘g’ (Lett et al., 2019), which is also in line with previously reported positive genetic correlations between CSA and EA in genomic (Grasby et al., 2020) and twin-studies (Vuoksiima et al., 2014). In the current study, we have further dissected the behavioural phenotype and assessed the mediation through sensitivity analyses, showing that the effect we observe is stronger for reading than for intelligence in the large sample of children from the ABCD study.

### Limitations

Despite performing a comprehensive investigation triangulating reading with the brain and its genetic effects, the current study also has some limitations.

First, the brain phenotypes included in the current study may not be the most adequate ones as we focused on the relationship between reading abilities and morphological cortical measures (CSA and CT). Although cortical morphometry can be seen as an indirect measure of function (Smith et al., 2019), functional task-related measures may be better phenotypes to establish the link from behaviour to the brain. Large datasets such as the ABCD and the UKB do not contain relevant reading-related task measures (Casey et al., 2018; Littlejohns et al., 2020), while smaller datasets do not provide enough power to perform the type of analyses carried out in the current study. However, functional connectivity from resting-state data, available in the ABCD and UKB, could be used as markers of interest in follow-up studies.

Second, we used a variety of analyses and datasets with different demographic characteristics. The goal was to find converging evidence across them, but this approach also adds heterogeneity that may hinder the interpretation of our results. We provide plausible explanations of why some results may not be congruent in terms of this heterogeneity, and, instead, focused on the strongest signals that replicated throughout the analyses to overcome this caveat.

Lastly, some crucial aspects to understand how reading affects the brain and its genetic underpinnings, such as development, were overlooked to some extent in the present study. As the ABCD is a longitudinal study that has been following children since 2018 and will continue to do so for the next 7 years, follow-up studies will have the opportunity to assess the stability of the effects reported here.

The present study provides a proof-of-concept of the type of the analytical approach that could be used to understand the biological bases of reading abilities, utilizing openly available datasets and tools. In the future, this approach could be extended to other relevant phenotypes, such as functional or structural connectivity measures. In addition, possible factors that are likely to shape the effects we describe (such as developmental age, or sex) should also be taken into consideration.

### Conclusion

In summary, we identified the cortical correlates of reading abilities, including total CSA and key reading-network measures such as the CSA of the STG and the CT of a cluster of occipital regions that are involved in the dorsal and ventral reading pathways. The effects reported in these analyses were robust and predominantly left-hemisphere specific. Nevertheless, these were also modest effects, which highlights the need for large datasets to be able to address these types of questions in an unbiased manner. Further, while there was no indication of a genetic overlap with any of the CT measures, we found evidence that suggests that the association with CSA measures is partly mediated by genetic effects. These findings revealed novel insights into the structural brain correlates of reading, and support the idea that genetics may contribute to defining the relation between reading with total CSA and superior temporal CSA measures in the left hemisphere.

## Methods

### ABCD study data

The Adolescent Brain Cognitive Development (ABCD) study is a longitudinal study across 21 data acquisition sites following 11,875 children starting at 9 and 10 years old (Jernigan & Brown, 2018). The study used specific recruitment strategies to create a population-based, demographically diverse sample (Garavan et al., 2018). However, it is not necessarily representative of the U.S. national population (Compton, Dowling, & Garavan, 2019).

The current study analysed the full baseline sample (N=11,878) from the ABCD data release 3.0 RDS (DOI: 10.15154/1520591) and the Genotyping Data from the ABCD Curated Annual Release 3.0 (NDA Study 901; DOI: 10.15154/1519007). All variables included in the current study are listed and described in Table S1, with Table S2 providing their descriptive statistics.

#### Behavioural data

The dependent variable for our primary analysis was the Toolbox Oral Reading Recognition Task from the NIH Toolbox Cognition Battery, which is a reading test that asks individuals to pronounce single words (Luciana et al., 2018). Three additional cognitive variables were included for the sensitivity analyses: Toolbox Picture Vocabulary Task (NIH Toolbox Cognition Battery), which measures language skills and verbal intellect; the “fluid composite” cognitive score, which is a composite score derived as the average normalized scores from several fluid ability measures from the NIH Toolbox Cognition Battery, namely: Flanker, Dimensional Change Card Sort, Picture Sequence Memory, List Sorting and Pattern Comparison; and the Matrix Reasoning task from the Wechsler Intelligence Scale for Children-V (WISC-V), which measures fluid intelligence and visuospatial reasoning.

#### Structural magnetic resonance imaging data

Destrieux (a2009s) Freesurfer atlas parcellation measures for left-hemisphere CT and CSA were included in the analysis. Seventy-four regional and one global measure (total CSA or mean CT) were included for CT and CSA. Thus, 150 measures were analyzed in total. The main analyses were focused on the left hemisphere, although some of the homotopic right-hemisphere measures were also included for completeness and sensitivity.

#### Additional data

The study comprises different data acquisition sites and associated MRI scanners. Therefore, the scanner machine id was included as a random factor to account for the dependence of samples, and family relatedness (family id) was included as a random factor nested within scanner in all analyses that included related individuals. The ABCD is a diverse dataset with a wide range of individuals across different ethnicities and socioeconomic backgrounds (Garavan et al., 2018). We included the following demographic variables in all analyses: sex, age, genetic principal components (PC1-PC10; see “Genetic analyses” section below), household income and parental education. Note that the genetic PCs were computed for the full sample (diverse ancestries) and for the European-PCA cluster for some genetic analyses, which aims to minimize variation in non-genetic factors and genetic factors (see below). The PCs corresponding to the specific dataset used in each analysis were included as covariates.

#### Brain-behaviour association analyses

##### Quality control

Individuals that passed quality control for neuroimaging (N=11,265) and genetic data (N=11,092) were selected. Next, we filtered out individuals that had missing data for covariates, or variables of interest (N=9,224), or with extreme outlier values (i.e. more than 7 standard deviations away from the mean for that variable: N=9,177). Finally, we also trimmed extreme values of the dependent variable (i.e. reading; see above) by excluding the 0.01 quantiles at each end of the distribution (N=164) to avoid having long tails that would affect the regression models (kurtosis was reduced from 4.56 to 3.43; see Figure S1). This final dataset of 9,013 individuals was used for the brain-behaviour association analyses. Descriptive statistics per covariate before and after the final trimming are presented in Table S2.

##### Regression analyses

All numeric variables were z-transformed (centered and scaled to have a mean of 0 and variance of 1). Linear mixed-effect regressions were run using R (v 4.0.3) package *lme4* (v 1.1-25; D. Bates et al., 2015) to identify structural ROIs associated with reading.

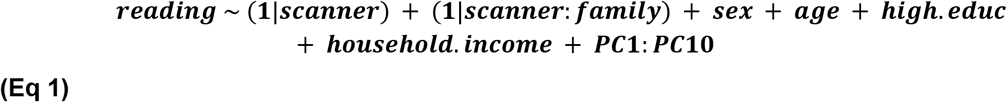

First, we included all random variables and covariates of interest in a baseline model (Equation 1; see Table S1 for the definition of all covariates and random factors). Each brain measure was then added as an independent variable in separate regressions to assess their effect on reading (Model 1). Next, we checked that the association of regional measures were robust after controlling for the corresponding global measures (Model 2), i.e. the left-hemisphere total CSA for CSA measures or the left-hemisphere mean CT for CT measures. A likelihood ratio test (LRT) between each pair of nested models (Baseline *vs* Model 1; Model 1 *vs* Model 2) was used to assess the significance of the term of interest. We defined brain measures robustly associated with reading as those that were significant after multiple testing corrections in both models 1 and 2 (False Discovery Rate q-value<0.05).

To assess whether the observed associations were specific to reading, we also explored whether the reading-associated measures were robust to adjustment for other cognitive covariates related to reading: fluid intelligence score and matrix reasoning, and picture vocabulary (see Table S1 for the definition of variables included in this sensitivity analysis).

Although the main analysis focused on left-hemisphere measures, the reading-associated measures were followed up by performing identical analyses with their homotopic right-hemisphere measures. We also computed the correlations between these homotopic regions and across all reading-associated brain measures, using the residuals after adjusting for the covariates in Models 1 and 2 (R package *psych* v2.0.12; Revelle, 2020).

We re-ran these analyses again restricting them to subsets of the data (see section below for exact subset definitions) to confirm that these results were not driven by the family component or were ancestry-specific: unrelated individuals only (after QC: N=7,502), and unrelated individuals of European ancestry (after QC: N=4,080).

All effects were visualized in brainplots using the R packages *ggseg* (v 1.6.4; Mowinckel & Piñero, 2020) and *ggsegDesterieux* (v 1.0.1.002; Mowinckel & Piñero, 2021).

#### Genetic analyses

##### Genetic quality control

Plink (v1.90b6.15; Purcell et al., 2007) was used to perform SNP and sample quality control (QC) following Coleman et al. (2015). 11,099 individuals had available genotype data for 516,598 SNPs. SNPs were excluded if they had genotyping rate <0.99, Hardy Weinberg Equilibrium (HWE) p-value <1e-6, or minor allele frequency (MAF) < 0.01. After this QC, 340,003 SNPs were kept. Samples were excluded if they had a missing genotyping rate greater than 95% (N=7).

A set of 291,223 common (MAF>0.05), autosomal, independent SNPs were selected by pruning LD using a window of 1500 variants and a shift of 150 variants between windows, with an r^2^ cut-off of 0.2, and excluding high-LD and non-autosomal regions (Coleman et al., 2015). These SNPs were used to identify sex mismatches, assess relatedness and flag outliers for IBD or heterozygosity, and to perform ancestry analyses. To identify a set of unrelated individuals, we randomly excluded one subject from each pair with a pi-hat > 0.1875. This resulted in 11,047 subjects passing genetic QC, of which 9,061 were unrelated (i.e. pi-hat>0.1875).

Genetic principal components (PCs) were calculated using SmartPCA (EIGENSOFT v 6.1.4; Patterson, Price, & Reich, 2006) for the total sample. These PCs were then used as covariates in the brain-behaviour regression analyses (see above). PCs computed with reference populations of the 1000 genomes reference dataset (v37; Auton et al., 2015) were plotted to visualize the genetic ancestry of the total sample.

In order to define a subset of homogeneous ancestry, we then selected individuals with a > 90% of European ancestry (as defined by the variable “genetic_ancestry_factor_european” from the ABCD data release 3.0; N=6,103) and removed outlier individuals exceeding 6 standard deviations along one of the top 10 PCs using SmartPCA (Patterson, Price, & Reich, 2006). The genetic PCs for this subset were used in downstream genetic analyses that are sensitive to population stratification effects. This subset consisted of 5,740 individuals of European ancestry, of which 4,716 were unrelated.

##### Heritability and genetic correlation analyses

A genetic relationship matrix (GRM) of 4,716 unrelated individuals of European ancestry was built in GCTA (v1.93.2beta; Yang et al., 2010) using 331,460 autosomal, directly genotyped SNPs. We further excluded one random participant from each pair having a kinship coefficient higher than 0.05 based on the calculated GRM (as this analysis is especially sensitive to higher levels of relatedness), resulting in 4,633 participants. Genome-based restricted maximum likelihood (GREML; Yang et al., 2010) analyses were performed to estimate the SNP-based heritabilities (h^2^-SNP), using residuals after controlling for the above-mentioned covariates and scanner (for brain measures) or site (for non-brain measures) as a random factor. The significance of the heritability estimates was Bonferroni corrected for multiple comparisons for the number of tested measures, i.e. 21 measures (9 brain measures, 8 brain measures adjusted for global measures and 4 cognitive measures). Bivariate GREML analyses were run to compute the genetic correlation (r_g_) between regional and global brain measures, and then assessed whether the r_g_ was significantly different to 1 through a one-tailed z-test (where z = (1-r_g_)/se).

### Heritability and genetic correlations based on summary statistics

LDSC was used to calculate the heritability and genetic correlations of the identified nine brain measures and three reading-related cognitive measures (as defined in Table S6). These included GWAS summary statistics for developmental dyslexia (Gialluisi et al., 2020), EA and cognitive performance (for fluid intelligence; Lee et al., 2018), and CT and CSA brain measures (Smith et al., 2021). The significance of the heritability estimates was Bonferroni corrected for multiple comparisons for the 12 tested measures.

### Polygenic Scoring

PRSice (Choi & OReilly, 2019) was used to compute polygenic scores (PGS) for each of the traits of interest (i.e. using the published cognitive GWAS summary statistics listed in Table S4 as base datasets).

The target dataset was the ABCD unrelated European ancestry subset (N=4,080). The ABCD imputed genotype data was used, which had been imputed using the TOPMed imputation server with Eagle v2.4 phasing and TOPMed mixed ancestry reference (http://dx.doi.org/10.15154/1519007). The imputed data was filtered using plink to keep only HRC biallelic genotype calls with a minimum genotype probability of 0.9, and SNPs with imputation quality scores above 0.7, and lifted to the hg19 reference panel (27,817,000 SNPs). Next, SNPs were filtered on HWE (p-value >1e-6), MAF>0.05 and missing genotype rate < 0.1 (4,034,417 SNPs kept).

The PGSs were then computed for each trait through clumping (clump-kb 250kb, clump-p: 1, clump-r2=0.1) and thresholding (8 p-value thresholds: 5e-08, 1e-5, 0.0001, 0.001, 0.05, 0.1, 0.5, 1). Next, linear mixed-effects models were run in R (v 4.0.3; package *lme4* v 1.1-25). A baseline model included the dependent variable and all covariates specified above (see section “Regression analyses”) and separate PGS models were run including each PGS as an independent variable. The significance of each PGS was assessed by a likelihood ratio test between the baseline and PGS model. The proportion of variance explained by the PGS (Δ*R*^2^) was computed as the difference of the *R*^2^ between the models.

The most predictive PGS for reading was first identified to define the most predictive base dataset and p-value threshold for reading in the ABCD dataset (see Figures S9, S10). Next, the selected base dataset and p-value threshold were used to perform the cross-trait regression (i.e. with the brain measures as dependent variables) in order to maximize power and limit the number of performed comparisons.

### Mediation analysis

For each model of interest (see Table 1), two regression models were computed using the R package *lme4* (Bates et al., 2015): an initial model that computed the effect of the independent variable (IV) on the mediator, and another that computed the indirect effect of the IV on the dependent variable, after accounting for the mediator. All models included the covariates and random structure indicated in Equation 1. The “mediation” package (Tingley et al., 2014)was then used to estimate causal mediation effects (Imai, Keele, & Tingley, 2010). The significance of the direct (ADE) and indirect (ACME) paths was assessed by nonparametric bootstrapping of the confidence intervals. Sensitivity analysis included additional covariates to assess if the mediation effects were robust to those adjustments (see Table 1).

## Data availability

ABCD data are publicly available through the National Institute of Mental Health (NIHM) Data Archive (https://data-archive.nimh.nih.gov/abcd). GWAS summary statistics used in this study are available from the NHGRI-EBI GWAS Catalog https://www.ebi.ac.uk/gwas/downloads/summary-statistics and the Oxford Brain Imaging Genetics Server – BIG40 (https://open.win.ox.ac.uk/ukbiobank/big40/).

## Code availability

The code associated to this study is publicly available at https://github.com/amaiacc/MS-brain-reading-genetics/.

## Acknowledgements

This research is supported by the Basque Government through the BERC 2022-2025 program and by the Spanish State Research Agency through BCBL Severo Ochoa excellence accreditation CEX2020-001010-S. A. C-C. is co-funded by the Spanish Ministry of Science and Innovation and the Agencia Estatal de Investigación through Ayudas Juan de la Cierva-Incorporación (ref. IJC2018-036023-I) and by the Programa Fellows Gipuzkoa de atracción y retención de talento from the Diputación Foral de Gipuzkoa. P. M. P-A. is supported by grants from the Spanish Ministry of Science and Innovation (PGC2018-093408-B-I00) and Neuroscience Research Projects programme from the Fundacion Tatiana Pérez de Guzmán el Bueno. M. C. is supported by the Spanish Ministry of Science and Innovation grant (RTI2018-093547-B-I00).

Data used in the preparation of this article were obtained from the ABCD Study (https://abcdstudy.org/) and are held in the NIMH Data Archive. This is a multisite, longitudinal study designed to recruit more than 10,000 children aged 9–10 and follow them over 10 years into early adulthood. The ABCD Study is supported by the National Institutes of Health (NIH) and additional federal partners under award numbers U01DA041022, U01DA041028, U01DA041048, U01DA041089, U01DA041106, U01DA041117, U01DA041120, U01DA041134, U01DA041148, U01DA041156, U01DA041174, U24DA041123, and U24DA041147. A full list of supporters is available at https://abcdstudy.org/federal-partners/. A listing of participating sites and a complete listing of the study investigators can be found at https://abcdstudy.org/principal-investigators/. ABCD consortium investigators designed and implemented the study and/or provided data but did not necessarily participate in the analysis or writing of this report. This manuscript reflects the views of the authors and may not reflect the opinions or views of the NIH or ABCD consortium investigators. The ABCD data repository grows and changes over time. The ABCD data used in this report came from https://doi.org/10.15154/1520591 and https://doi.org/10.15154/1519007 (genotyping data).

## Notes

### Competing Interest Statement

The authors have declared no competing interest.

## References

Alfaro-Almagro, F., McCarthy, P., Afyouni, S., Andersson, J. L. R., Bastiani, M., Miller, K. L., … Smith, S. M. (2021). Confound modelling in UK biobank brain imaging. NeuroImage, 224, 117002. https://doi.org/10.1016/j.neuroimage.2020.117002

Andreola, C., Mascheretti, S., Belotti, R., Ogliari, A., Marino, C., Battaglia, M., & Scaini, S. (2021). The heritability of reading and reading-related neurocognitive components: A multi-level meta-analysis. Neuroscience & Biobehavioral Reviews, 121, 175–200. https://doi.org/10.1016/j.neubiorev.2020.11.016

Atteveldt, N. van, Formisano, E., Goebel, R., & Blomert, L. (2004). Integration of letters and speech sounds in the human brain. Neuron, 43(2), 271–282. https://doi.org/10.1016/j.neuron.2004.06.025

Auton, A., Abecasis, G. R., Altshuler, D. M., Durbin, R. M., Abecasis, G. R., Bentley, D. R., … and. (2015). A global reference for human genetic variation. Nature, 526(7571), 68–74. https://doi.org/10.1038/nature15393

Bates, D., Mächler, M., Bolker, B., & Walker, S. (2015). Fitting linear mixed-effects models using lme4. Journal of Statistical Software, 67(1), 1–48. https://doi.org/10.18637/jss.v067.i01

Ben-Shachar, M., Dougherty, R. F., & Wandell, B. A. (2007). White matter pathways in reading. Current Opinion in Neurobiology, 17(2), 258–270. https://doi.org/10.1016/j.conb.2007.03.006

Boets, B., Beeck, H. P. O. de, Vandermosten, M., Scott, S. K., Gillebert, C. R., Mantini, D., … Ghesquiere, P. (2013). Intact but less accessible phonetic representations in adults with dyslexia. Science, 342(6163), 1251–1254. https://doi.org/10.1126/science.1244333

Bulik-Sullivan, B., Finucane, H. K., Anttila, V., Gusev, A., Day, F. R., Loh, P.-R., … Neale, B. M. (2015). An atlas of genetic correlations across human diseases and traits. Nature Genetics, 47(11), 1236–1241. https://doi.org/10.1038/ng.3406

Carreiras, M., Quiñones, I., Hernández-Cabrera, J. A., & Duñabeitia, J. A. (2015). Orthographic coding: Brain activation for letters, symbols, and digits. Cerebral Cortex, 25(12), 4748–4760. https://doi.org/10.1093/cercor/bhu163

Casey, B. J., Cannonier, T., Conley, M. I., Cohen, A. O., Barch, D. M., Heitzeg, M. M., … Dale, A. M. (2018). The adolescent brain cognitive development (ABCD) study: Imaging acquisition across 21 sites. Developmental Cognitive Neuroscience, 32, 43–54. https://doi.org/10.1016/j.dcn.2018.03.001

Choi, S. W., & OReilly, P. F. (2019). PRSice-2: Polygenic risk score software for biobank-scale data. GigaScience, 8(7). https://doi.org/10.1093/gigascience/giz082

Cohen, L., Dehaene, S., Naccache, L., Lehéricy, S., Dehaene-Lambertz, G., Hénaff, M.-A., & Michel, F. (2000). The visual word form area. Brain, 123(2), 291–307. https://doi.org/10.1093/brain/123.2.291

Coleman, J. R. I., Euesden, J., Patel, H., Folarin, A. A., Newhouse, S., & Breen, G. (2015). Quality control, imputation and analysis of genome-wide genotyping data from the illumina HumanCoreExome microarray. Briefings in Functional Genomics, 15(4), 298–304. https://doi.org/10.1093/bfgp/elv037

Compton, W. M., Dowling, G. J., & Garavan, H. (2019). Ensuring the best use of data. JAMA Pediatrics, 173(9), 809. https://doi.org/10.1001/jamapediatrics.2019.2081

Dehaene, S., & Cohen, L. (2007). Cultural recycling of cortical maps. Neuron, 56(2), 384–398. https://doi.org/10.1016/j.neuron.2007.10.004

Dehaene, S., Cohen, L., Morais, J., & Kolinsky, R. (2015). Illiterate to literate: Behavioural and cerebral changes induced by reading acquisition. Nature Reviews Neuroscience, 16(4), 234–244. https://doi.org/10.1038/nrn3924

Destrieux, C., Fischl, B., Dale, A., & Halgren, E. (2010). Automatic parcellation of human cortical gyri and sulci using standard anatomical nomenclature. NeuroImage, 53(1), 1–15. https://doi.org/10.1016/j.neuroimage.2010.06.010

Dick, A. S., Lopez, D. A., Watts, A. L., Heeringa, S., Reuter, C., Bartsch, H., … Thompson, W. K. (2021). Meaningful associations in the adolescent brain cognitive development study. NeuroImage, 239, 118262. https://doi.org/10.1016/j.neuroimage.2021.118262

Doust, C., Fontanillas, P., Eising, E., Gordon, S. D., Wang, Z., Alagöz, G., … Luciano, M. (2021). Discovery of 42 genome-wide significant loci associated with dyslexia. medRxiv. https://doi.org/10.1101/2021.08.20.21262334

Eising, E., Mirza-Schreiber, N., Zeeuw, E. L. de, Wang, C. A., Truong, D. T., Allegrini, A. G., … Fisher, S. E. (2021). Genome-wide association analyses of individual differences in quantitatively assessed reading- and language-related skills in up to 34,000 people. bioRxiv. https://doi.org/10.1101/2021.11.04.466897

Elliott, L. T., Sharp, K., Alfaro-Almagro, F., Shi, S., Miller, K. L., Douaud, G., … Smith, S. M. (2018). Genome-wide association studies of brain imaging phenotypes in UK biobank. Nature, 562(7726), 210–216. https://doi.org/10.1038/s41586-018-0571-7

Feng, X., Altarelli, I., Monzalvo, K., Ding, G., Ramus, F., Shu, H., … Dehaene-Lambertz, G. (2020). A universal reading network and its modulation by writing system and reading ability in French and Chinese children. eLife, 9. https://doi.org/10.7554/elife.54591

Fisher, S. E., & DeFries, J. C. (2002). Developmental dyslexia: Genetic dissection of a complex cognitive trait. Nature Reviews Neuroscience, 3(10), 767–780. https://doi.org/10.1038/nrn936

Frei, O., Holland, D., Smeland, O. B., Shadrin, A. A., Fan, C. C., Maeland, S., … Dale, A. M. (2019). Bivariate causal mixture model quantifies polygenic overlap between complex traits beyond genetic correlation. Nature Communications, 10(1). https://doi.org/10.1038/s41467-019-10310-0

Friederici, A. D. (2012). The cortical language circuit: From auditory perception to sentence comprehension. Trends in Cognitive Sciences, 16(5), 262–268. https://doi.org/10.1016/j.tics.2012.04.001

Garavan, H., Bartsch, H., Conway, K., Decastro, A., Goldstein, R. Z., Heeringa, S., … Zahs, D. (2018). Recruiting the ABCD sample: Design considerations and procedures. Developmental Cognitive Neuroscience, 32, 16–22. https://doi.org/10.1016/j.dcn.2018.04.004

Gialluisi, A., Andlauer, T. F. M., Mirza-Schreiber, N., Moll, K., Becker, J., Hoffmann, P., … Schulte-Körne, G. (2020). Genome-wide association study reveals new insights into the heritability and genetic correlates of developmental dyslexia. Molecular Psychiatry. https://doi.org/10.1038/s4138002000898x

Grasby, K. L., Jahanshad, N., Painter, J. N., Colodro-Conde, L., Bralten, J., Hibar, D. P., … Mari, Z. (2020). The genetic architecture of the human cerebral cortex. Science, 367(6484).

Hickok, G., & Poeppel, D. (2007). The cortical organization of speech processing. Nature Reviews Neuroscience, 8(5), 393–402. https://doi.org/10.1038/nrn2113

Hyatt, C. S., Owens, M. M., Crowe, M. L., Carter, N. T., Lynam, D. R., & Miller, J. D. (2020). The quandary of covarying: A brief review and empirical examination of covariate use in structural neuroimaging studies on psychological variables. NeuroImage, 205, 116225. https://doi.org/10.1016/j.neuroimage.2019.116225

Imai, K., Keele, L., & Tingley, D. (2010). A general approach to causal mediation analysis. Psychological Methods, 15(4), 309–334. https://doi.org/10.1037/a0020761

Jernigan, T. L., & Brown, S. A. (2018). Introduction. Developmental Cognitive Neuroscience, 32, 1–3. https://doi.org/10.1016/j.dcn.2018.02.002

Landi, N., & Perdue, M. V. (2019). Neuroimaging genetics studies of specific reading disability and developmental language disorder: A review. Language and Linguistics Compass, 13(9). https://doi.org/10.1111/lnc3.12349

Lau, E. F., Phillips, C., & Poeppel, D. (2008). A cortical network for semantics: (De)constructing the N400. Nature Reviews Neuroscience, 9(12), 920–933. https://doi.org/10.1038/nrn2532

Lee, J. J., Wedow, R., Okbay, A., Kong, E., Maghzian, O., Zacher, M., … Cesarini, D. (2018). Gene discovery and polygenic prediction from a genome-wide association study of educational attainment in 1.1 million individuals. Nature Genetics, 50(8), 1112–1121. https://doi.org/10.1038/s41588-018-0147-3

Lett, T. A., Vogel, B. O., Ripke, S., Wackerhagen, C., Erk, S., Awasthi, S., … and, H. W. (2019). Cortical surfaces mediate the relationship between polygenic scores for intelligence and general intelligence. Cerebral Cortex, 30(4), 2708–2719. https://doi.org/10.1093/cercor/bhz270

Littlejohns, T. J., Holliday, J., Gibson, L. M., Garratt, S., Oesingmann, N., Alfaro-Almagro, F., … Allen, N. E. (2020). The UK biobank imaging enhancement of 100,000 participants:0.167emrationale, data collection, management and future directions. Nature Communications, 11(1). https://doi.org/10.1038/s41467-020-15948-9

Luciana, M., Bjork, J. M., Nagel, B. J., Barch, D. M., Gonzalez, R., Nixon, S. J., & Banich, M. T. (2018). Adolescent neurocognitive development and impacts of substance use: Overview of the adolescent brain cognitive development (ABCD) baseline neurocognition battery. Developmental Cognitive Neuroscience, 32, 67–79. https://doi.org/10.1016/j.dcn.2018.02.006

Maier, R. M., Visscher, P. M., Robinson, M. R., & Wray, N. R. (2018). Embracing polygenicity: A review of methods and tools for psychiatric genetics research. Psychological Medicine, 48(7), 1055–1067. https://doi.org/10.1017/s0033291717002318

Martin, A., Schurz, M., Kronbichler, M., & Richlan, F. (2015). Reading in the brain of children and adults: A meta-analysis of 40 functional magnetic resonance imaging studies. Human Brain Mapping, 36(5), 1963–1981. https://doi.org/10.1002/hbm.22749

Mitchell, B. L., Cuéllar-Partida, G., Grasby, K. L., Campos, A. I., Strike, L. T., Hwang, L.-D., … Rentería, M. E. (2020). Educational attainment polygenic scores are associated with cortical total surface area and regions important for language and memory. NeuroImage, 116691. https://doi.org/10.1016/j.neuroimage.2020.116691

Mowinckel, Athanasia M., & Vidal-Piñeiro, D. (2020). Visualization of brain statistics with r packages ggseg and ggseg3d. Advances in Methods and Practices in Psychological Science, 3(4), 466–483. https://doi.org/10.1177/2515245920928009

Mowinckel, Athanasia Mo, & Vidal-Piñeiro, D. (2021). ggsegDesterieux: Desterieux datasets for the ggseg-plotting tool. Retrieved from https://github.com/LCBC-UiO/ggsegDesterieux

Palmer, C. E., Zhao, W., Loughnan, R., Fan, C. C., Thompson, W., Jernigan, T. L., & Dale, A. M. (2019). Determining the association between cortical morphology and cognition in 10,145 children from the adolescent brain and cognitive development (ABCD) study using the MOSTest. bioRxiv. https://doi.org/10.1101/816025

Patterson, N., Price, A. L., & Reich, D. (2006). Population structure and eigenanalysis. PLoS Genetics, 2(12), e190. https://doi.org/10.1371/journal.pgen.0020190

Perdue, M. V., Mednick, J., Pugh, K. R., & Landi, N. (2020). Gray matter structure is associated with reading skill in typically developing young readers. Cerebral Cortex, 30(10), 5449–5459. https://doi.org/10.1093/cercor/bhaa126

Pugh, K. R., Mencl, W. E., Jenner, A. R., Katz, L., Frost, S. J., Lee, J. R., … Shaywitz, B. A. (2001). Neurobiological studies of reading and reading disability. Journal of Communication Disorders, 34(6), 479–492. https://doi.org/10.1016/s0021-9924(01)00060-0

Purcell, S., Neale, B., Todd-Brown, K., Thomas, L., Ferreira, M. A. R., Bender, D., … Sham, P. C. (2007). PLINK: A tool set for whole-genome association and population-based linkage analyses. The American Journal of Human Genetics, 81(3), 559–575. https://doi.org/10.1086/519795

Płoński, P., Gradkowski, W., Altarelli, I., Monzalvo, K., Ermingen-Marbach, M. van, Grande, M., … Jednoróg, K. (2016). Multi-parameter machine learning approach to the neuroanatomical basis of developmental dyslexia. Human Brain Mapping, 38(2), 900–908. https://doi.org/10.1002/hbm.23426

Revelle, W. (2020). Psych: Procedures for psychological, psychometric, and personality research. Evanston, Illinois: Northwestern University. Retrieved from https://CRAN.R-project.org/package=psych

Roeske, D., Ludwig, K. U., Neuhoff, N., Becker, J., Bartling, J., Bruder, J., … Schulte-Körne, G. (2009). First genome-wide association scan on neurophysiological endophenotypes points to trans-regulation effects on SLC2A3 in dyslexic children. Molecular Psychiatry, 16(1), 97–107. https://doi.org/10.1038/mp.2009.102

Rueckl, J. G., Paz-Alonso, P. M., Molfese, P. J., Kuo, W.-J., Bick, A., Frost, S. J., … Frost, R. (2015). Universal brain signature of proficient reading: Evidence from four contrasting languages. Proceedings of the National Academy of Sciences, 112(50), 15510–15515. https://doi.org/10.1073/pnas.1509321112

Sandak, R., Mencl, W. E., Frost, S. J., & Pugh, K. R. (2004). The neurobiological basis of skilled and impaired reading: Recent findings and new directions. Scientific Studies of Reading, 8(3), 273–292. https://doi.org/10.1207/s1532799xssr0803_6

Smith, S., Duff, E., Groves, A., Nichols, T. E., Jbabdi, S., Westlye, L. T., … Douaud, G. (2019). Structural variability in the human brain reflects fine-grained functional architecture at the population level. The Journal of Neuroscience, 39(31), 6136–6149. https://doi.org/10.1523/jneurosci.2912-18.2019

Smith, S. M., Douaud, G., Chen, W., Hanayik, T., Alfaro-Almagro, F., Sharp, K., & Elliott, L. T. (2021). An expanded set of genome-wide association studies of brain imaging phenotypes in UK biobank. Nature Neuroscience. https://doi.org/10.1038/s41593-021-00826-4

Tingley, D., Yamamoto, T., Hirose, K., Keele, L., & Imai, K. (2014). Mediation:RPackage for causal mediation analysis. Journal of Statistical Software, 59(5). https://doi.org/10.18637/jss.v059.i05

Verhoef, E., Shapland, C. Y., Fisher, S. E., Dale, P. S., & Pourcain, B. S. (2020). The developmental origins of genetic factors influencing language and literacy: Associations with early-childhood vocabulary. Journal of Child Psychology and Psychiatry. https://doi.org/10.1111/jcpp.13327

Visscher, P. M., Hemani, G., Vinkhuyzen, A. A. E., Chen, G.-B., Lee, S. H., Wray, N. R., … Yang, J. (2014). Statistical power to detect genetic (co)variance of complex traits using SNP data in unrelated samples. PLoS Genetics, 10(4), e1004269. https://doi.org/10.1371/journal.pgen.1004269

Yang, J., Benyamin, B., McEvoy, B. P., Gordon, S., Henders, A. K., Nyholt, D. R., … Visscher, P. M. (2010). Common SNPs explain a large proportion of the heritability for human height. Nat. Genet., 42(7), 565–569.

