## Supplementary information for "Brain structure, phenotypic and genetic correlates of reading abilities"

##### List of supplementary tables

**Table S1: Data tables and variables (ABCD 3.0 release RDS)**.

**Table S2: Descriptive statistics of the ABCD data for variables used in the brain-behaviour association analysis**.

**Table S3: Effects of covariates on the baseline model for reading.**

**Table S4: Brain-reading association results.**

**Table S5: Sensitivity analysis for the reading-associated brain measure.**

**Table S6: Sensitivity analyses of the brain-behaviour associations after adjusting for additional cognitive covariates**.

**Table S7: Publicly available GWAS summary statistics used in the current study**.

**Table S8: Heritability estimates for reading and related traits, across methods**.

#### Supplementary figures

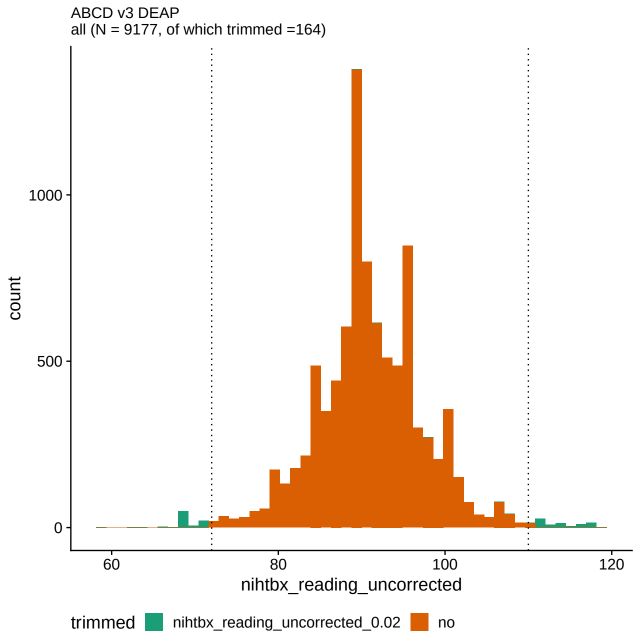


**Figure S1:** Distribution of reading in the ABCD baseline dataset. The 1% quantiles at each end of the distribution are indicated with the dashed vertical lines, and the excluded individuals are coloured in green (i.e. trimmed individuals).

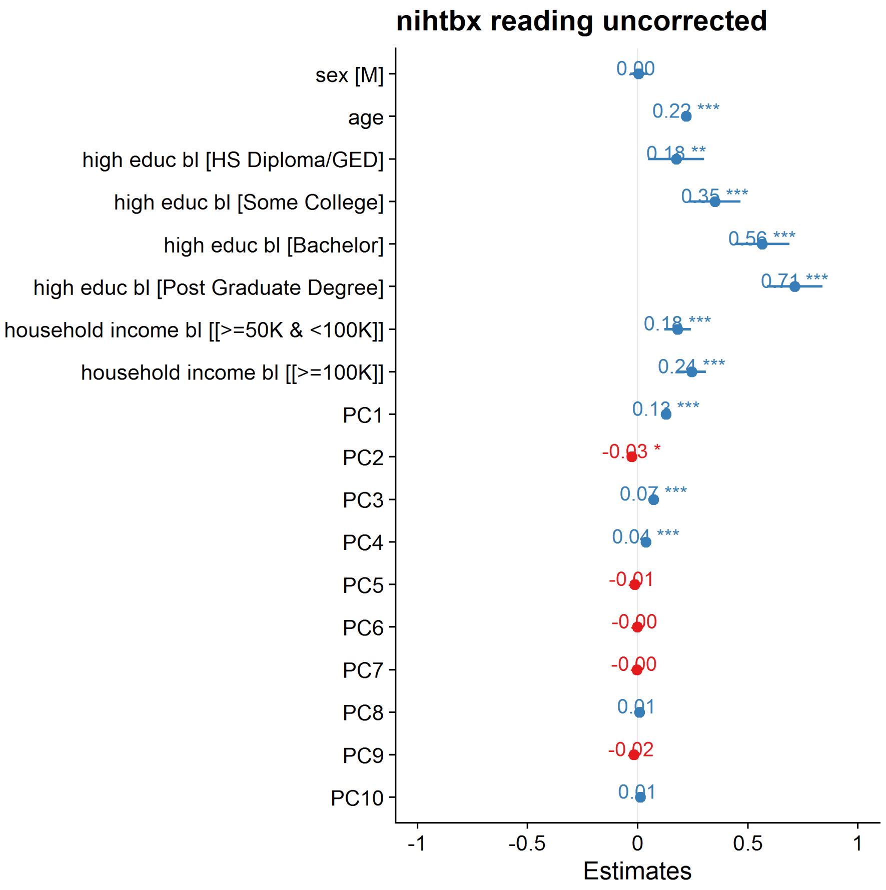


**Figure S2:** Estimates for the baseline regression model: $reading \sim covariates$.

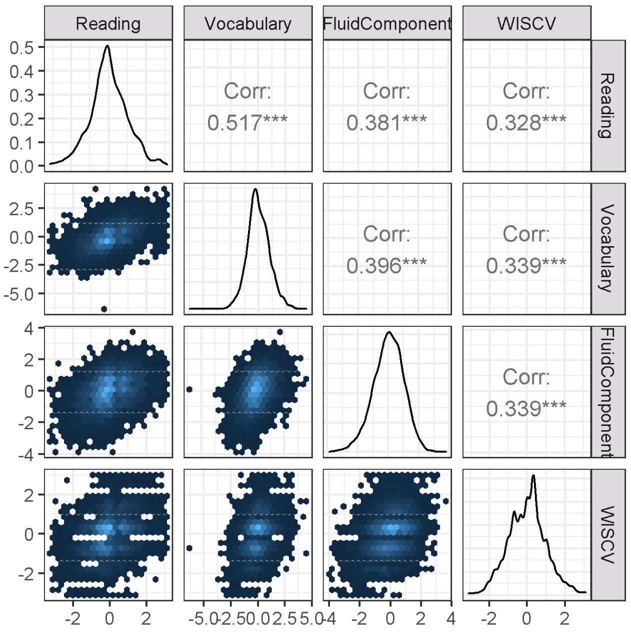


**Figure S3: Distributions and phenotypic correlations across reading and cognitive measures (ABCD dataset; total sample N=9,013).** The diagonal shows the distribution of the standardized trait, the upper triangle shows correlation coefficients, and the lower triangle shows the scatterplot for each pair of variables (colour representing the density of data-points within that bin). FluidComponent = NIHTBX fluid component; Vocabulary = NIHTBX picture vocabulary; WISCV = WISC-V Matrix Reasoning Total Scaled Score

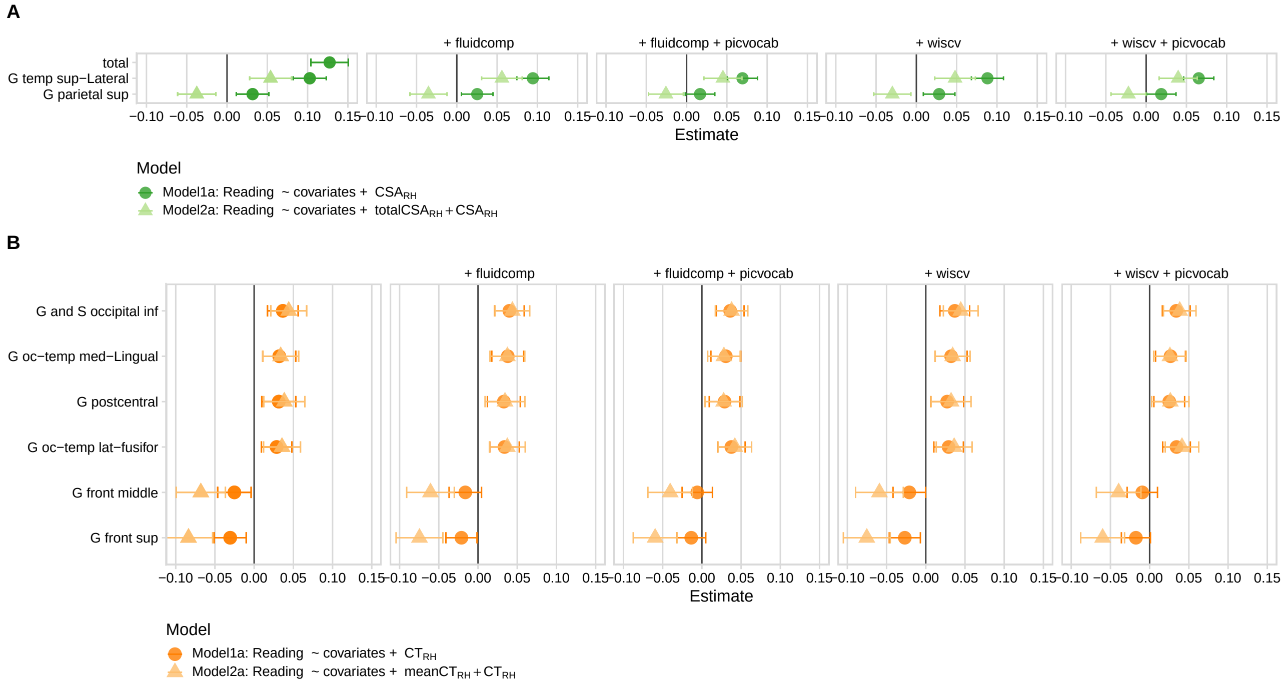


**Figure S4:** Sensitivity analysis of the effect of reading-associated measures on reading ability. Beta estimates for **A** cortical surface area measures and **B** cortical thickness measures after adjusting for additional cognitive variables. Model1: test model, Model2: test model after adjusting for global measure (total CSA or mean CT). Fluidcomp= NIHTBX fluid component; Picvocab= NIHTBX picture vocabulary; WISCV = WISC-V Matrix Reasoning Total Scaled Score. The significant regions are highlighted in bold.

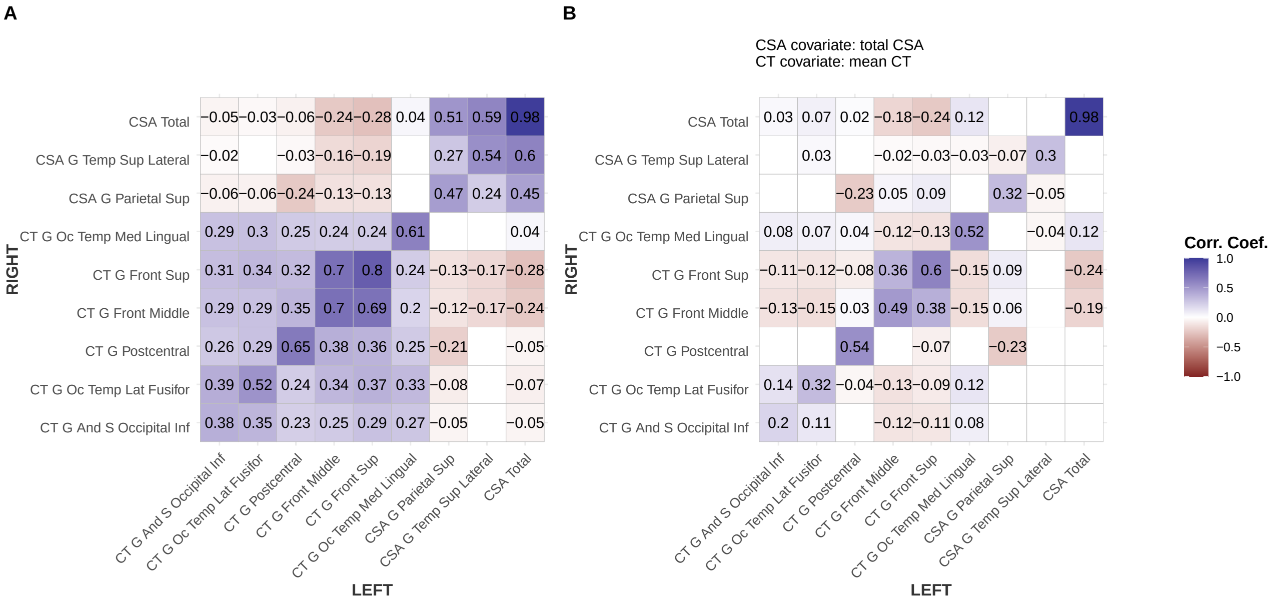


**Figure S5:** Phenotypic correlations across brain measures associated with reading: (**A**) after adjusting for covariates and (**B**) after adjusting for covariates and global brain measures. The diagonal shows the left-right correlation for each measure. The upper triangle shows correlations in right hemisphere measures and the lower triangle shows correlations across left hemisphere measures. Non-significant correlations (Bonferroni adjusted) are blanked.

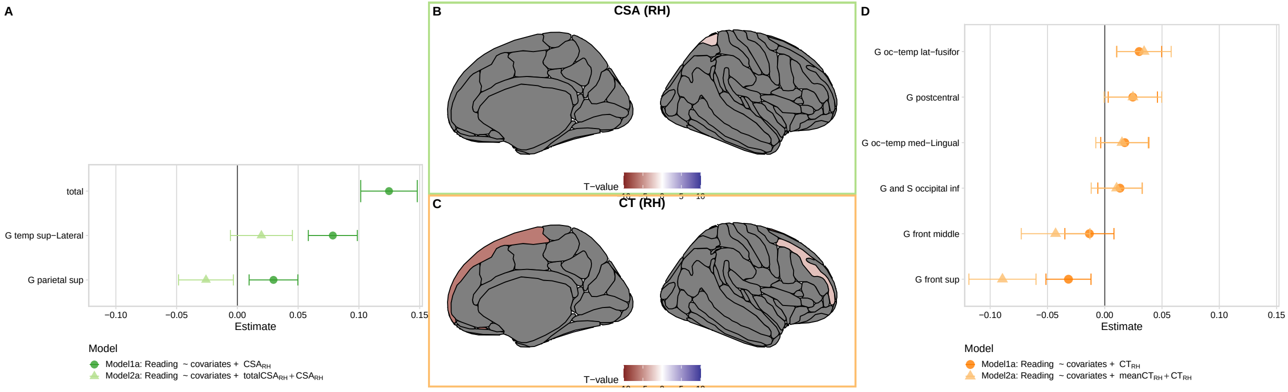


**Figure S6:** Effect of reading-associated homotopic measures (right-hemisphere) on reading ability. Beta estimates for **A** cortical surface area measures and **B** cortical thickness measures. Model1: test model, Model2: test model after adjusting for global measure (total CSA or mean CT). The significant regions are highlighted in bold.

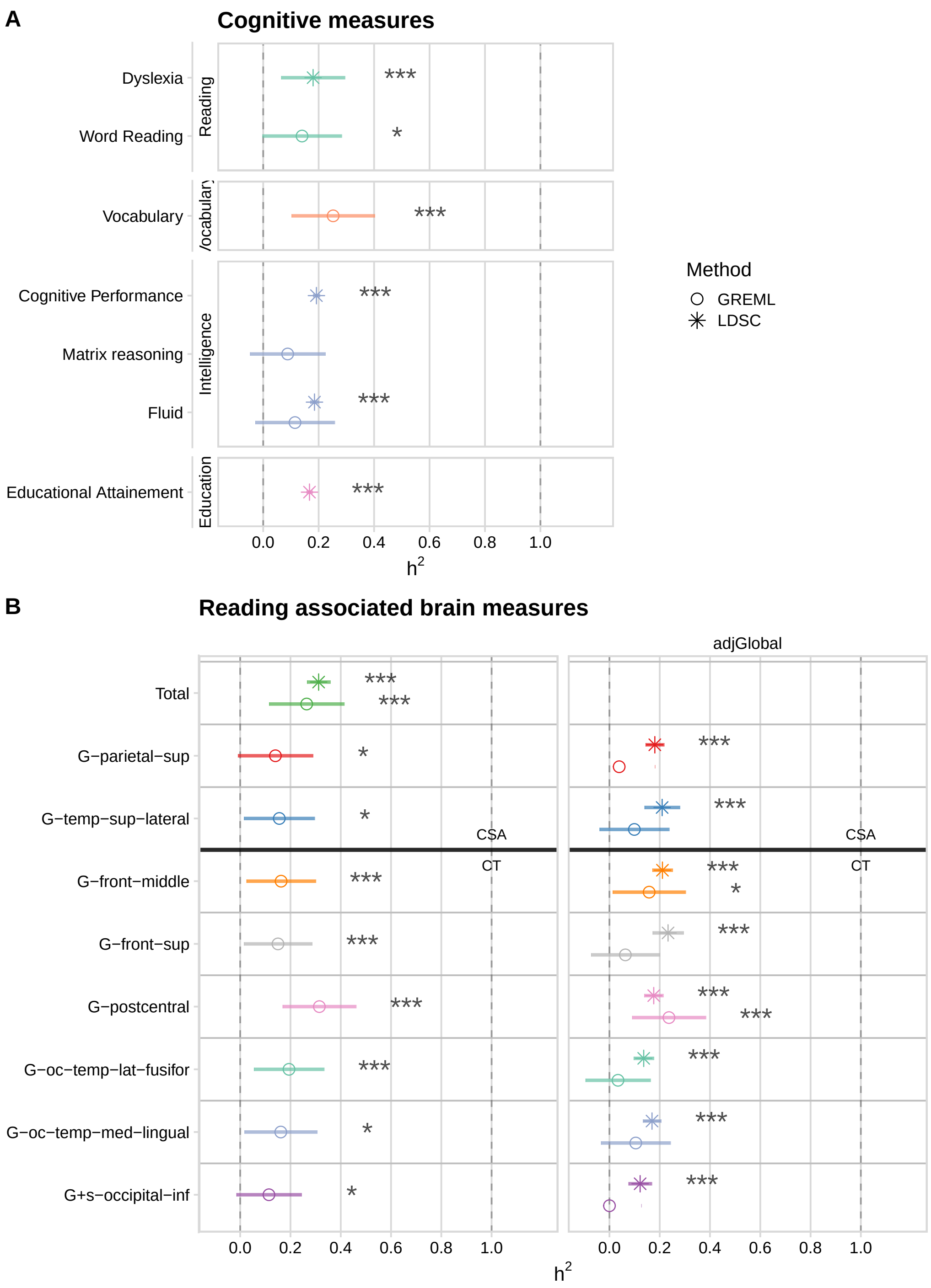


**Figure S7:** Heritability for (**A**) reading and related cognitive measures and (**B**) reading-associated brain measures. Points represent the estimate and the error bars indicate the 95% confidence interval. CSA = cortical surface area; CT = cortical thickness; G = gyrus; S = sulcus; GREML = genome-based restricted maximum likelihood; LDSC = linkage disequilibrium score regression; adjGlobal= estimates after adjusting for global brain measures; the global measure was 'head size' for LDSC-based estimates from the UK Biobank (Smith et al., 2021); GREML estimates based on the ABCD data: global measures were 'total left CSA' for CSA measures, 'mean left CT' for CT measures. * = uncorrected p-value $< 0.05$; *** = p-value Bonferroni adjusted for the number of cognitive measures (8) or brain measures (9) $< 0.05$.

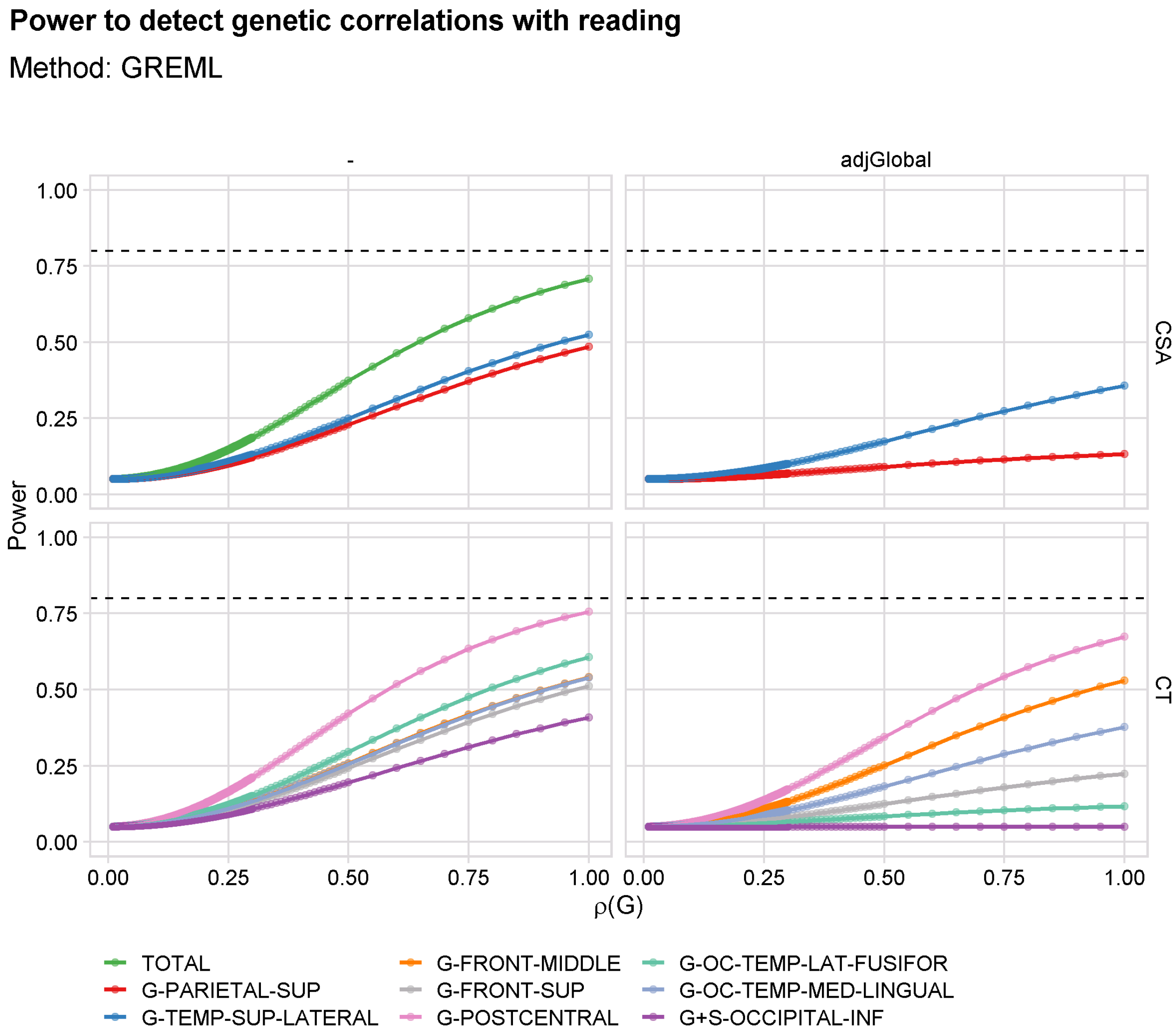


**Figure S8:** Approximate power to detect genetic correlation ($\rho$(G)) between reading and reading-associated brain measures using bivariate GREML in the ABCD unrelated European-PCA dataset (N=4,633). The GCTA power calculator (Visscher et al., 2014)was used in all cases, using the SNP-based heritability estimates for each trait ($h^2_{SNP}$). See Figure S7 for the $h^2_{SNP}$ estimates. The dashed horizontal line indicates $\beta$=0.80.

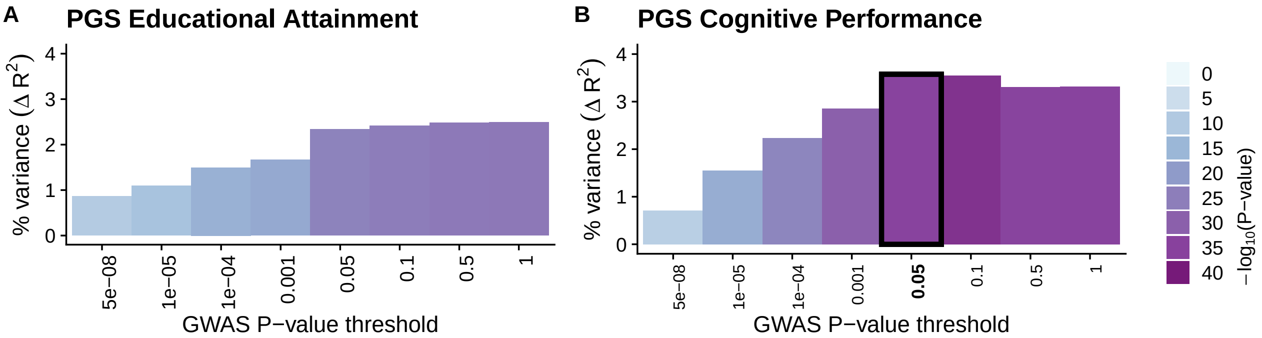


**Figure S9: Prediction results for reading using PGS**. PGS for (**A**) educational attainment and (**B**) cognitive performance. Summary statistics from Lee et al., 2018. The best PGS for reading is highlighted in bold.

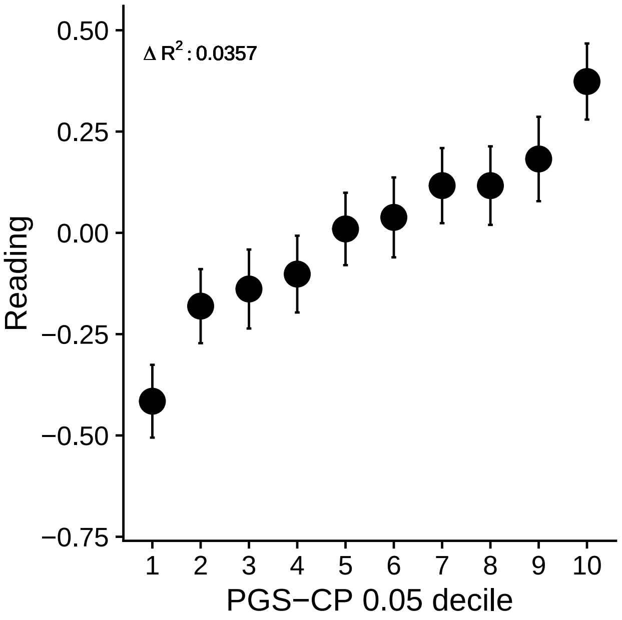


**Figure S10: Decile plot for PGS of CP on reading**.

### References

Lee, J. J., Wedow, R., Okbay, A., Kong, E., Maghzian, O., Zacher, M., Nguyen-Viet, T. A., Bowers, P., Sidorenko, J., Linnér, R. K., Fontana, M. A., Kundu, T., Lee, C., Li, H., Li, R., Royer, R., Timshel, P. N., Walters, R. K., Willoughby, E. A., ... Cesarini, D. (2018). Gene discovery and polygenic prediction from a genome-wide association study of educational attainment in 1.1 million individuals. *Nature Genetics*, *50*(8), 1112–1121.

Smith, S. M., Douaud, G., Chen, W., Hanayik, T., Alfaro-Almagro, F., Sharp, K., & Elliott, L. T. (2021). An expanded set of genome-wide association studies of brain imaging phenotypes in UK biobank. *Nature Neuroscience*.

Visscher, P. M., Hemani, G., Vinkhuyzen, A. A. E., Chen, G.-B., Lee, S. H., Wray, N. R., Goddard, M. E., & Yang, J. (2014). Statistical power to detect genetic (co)variance of complex traits using SNP data in unrelated samples (G. S. Barsh, Ed.). *PLoS Genetics*, *10*(4), e1004269.
